# Palmitoylation of gasdermin D directs its membrane translocation and pore formation in pyroptosis

**DOI:** 10.1101/2023.02.21.529402

**Authors:** Arumugam Balasubramanian, Laxman Ghimire, Alan Y. Hsu, Hiroto Kambara, Xing Liu, Tomoya Hasegawa, Rong Xu, Muhammad Tahir, Hongbo Yu, Judy Lieberman, Hongbo R. Luo

**Affiliations:** Department of Pathology, Dana-Farber/Harvard Cancer Center, Harvard Medical School; Department of Laboratory Medicine, Boston Children’s Hospital, Enders Research Building, Room 811, Boston, MA, 02115, USA; Program in Cellular and Molecular Medicine, Boston Children’s Hospital, Boston, MA 02115, USA; VA Boston Healthcare System, Department of Pathology and Laboratory Medicine, 1400 VFW Parkway, West Roxbury, MA 02132 USA; Department of Pediatrics, Harvard Medical School, Boston, MA 02115, USA

## Abstract

Gasdermin D (GSDMD)-mediated macrophage pyroptosis plays a critical role in inflammation and host defense. Plasma membrane perforation elicited by caspase-cleaved GSDMD N-terminal domain (GSDMD-NT) triggers membrane rupture and subsequent pyroptotic cell death, resulting in release of pro-inflammatory IL-1β and IL-18. However, the biological processes leading to its membrane translocation and pore formation are not fully understood. Here, using a proteomics approach, we identified fatty acid synthase (FASN) as a GSDMD-binding partner and demonstrated that post-translational palmitoylation of GSDMD at Cys191/Cys192 (human/mouse) led to membrane translocation of GSDMD-NT but not full-length GSDMD. GSDMD lipidation, mediated by palmitoyl acyltransferases ZDHHC5/9 and facilitated by LPS-induced reactive oxygen species (ROS), was essential for GSDMD pore-forming activity and pyroptosis. Inhibition of GSDMD palmitoylation with palmitate analog 2-bromopalmitate or a cell permeable GSDMD-specific competing peptide suppressed pyroptosis and IL-1β release in macrophages, mitigated organ damage, and extended the survival of septic mice. Collectively, we establish GSDMD-NT palmitoylation as a key regulatory mechanism controlling GSDMD membrane localization and activation, providing a novel target for modulating immune activity in infectious and inflammatory diseases.

**One Sentence Summary:** LPS-induced palmitoylation at Cys191/Cys192 is required for GSDMD membrane translocation and its pore-forming activity in macrophages.

## Introduction

Pyroptosis, a form of lytic cell death triggered by proinflammatory signals, plays a critical role in inflammation and host defense responses(*1, 2*). The pore-forming protein gasdermin D (GSDMD) is an executor of pyroptosis(*3–6*). Upon activation of the inflammasome, caspase-1 cleaves GSDMD into a GSDMD N-terminal pore-forming domain and a C-terminal autoinhibitory domain(*3, 7*). The cleaved N-terminal domain (GSDMD-NT) binds to phosphatidylinositol phosphates and phosphatidylserine in the cell membrane inner leaflet to form large oligomers, which generate approximately 20 nm pores in the plasma membrane and consequent lytic cell death(*3, 5, 8–15*). GSDMD can also be cleaved and activated by LPS-induced activation of non-canonical inflammasomes via murine caspase-11 (or human caspase-4 and −5). Additionally, caspase-8 and neutrophil elastase (ELANE) cleave GSDMD to induce lytic cell death in certain contexts(*16–19*). GSDMD-NT-mediated plasma membrane perforation triggers profound membrane rupture and subsequent lytic death to ultimately facilitate clearance of intracellular pathogens and the maturation and release of proinflammatory cytokines such as IL-1β and IL-18(*3–5, 15, 20*). A recent study suggested that GSDMD also plays an essential role in inflammasome-mediated IL-1β secretion from living macrophages(*20*).

GSDMD cleavage and GSDMD-mediated lytic cell death are tightly regulated. For instance, succination of GSDMD Cys191/Cys192 (human/mouse) prevents GSDMD-caspase interactions, limiting GSDMD processing, oligomerization, and its ability to induce cell death(*21*). The Ragulator-Rag-mTORCI pathway, by mediating reactive oxygen species (ROS) production, promotes GSDMD oligomerization but not its membrane localization, providing a checkpoint for pore formation and consequent pyroptosis and uncoupling biochemical GSDMD cleavage from pore formation(*22*). Oligomerization of GSDMD-NT at the plasma membrane produces a specialized plasma membrane pore structure that preferentially releases mature IL-1β and non-selective ionic fluxes(*15*). Oncotic cell swelling then ruptures the plasma membrane, which is not simply a passive event mediated by osmotic lysis but tightly regulated by cell surface NINJ1 protein during lytic pyroptotic cell death(*23*). A recent study also suggested that GSDMD pores are dynamic structures, with pore-mediated calcium influx modulating pore opening/closing kinetics through phosphoinositide metabolism(*24*). However, whether GSDMD plasma membrane translocation after its cleavage is also tightly controlled and how this process is regulated remain elusive.

### GSDMD interacts with fatty acid synthase (FASN) and can be palmitoylated

To further elucidate the physiological mechanisms controlling GSDMD processing and activation, we performed a proteomics analysis to identify GSDMD-interacting proteins. We expressed full length GSDMD (GSDMD-FL) in HEK293T cells and immunoprecipitated GSDMD-containing protein complexes using a specific antibody targeting GSDMD (**fig. S1a**). Co-immunoprecipitated proteins were eluted and analyzed by liquid chromatography-tandem mass spectrometry (LC-MS/MS) (**Table S1**). Proteomics analysis identified FASN as a GSDMD-interacting protein (**Table S2**). Western blotting confirmed endogenous FASN expression in HEK293T cells and its co-immunoprecipitation with recombinant GSDMD but not actin or control antibodies (**fig. S1b**). To investigate whether endogenous GSDMD and FASN interact under pathophysiological conditions, PMA-differentiated macrophage-like THP-1 cells were treated with or without LPS. Co-IP analysis revealed that FASN interacted with GSDMD only in the presence of LPS (**fig. S1c**), suggesting that this interaction requires an LPS-elicited event in macrophages.

FASN is responsible for the catalysis of carbohydrate-derived glycolytic precursors into long-chain fatty acids. It also catalyzes palmitate biosynthesis and is thus required for protein palmitoylation, a post-translational modification (PTM) where palmitic acid is covalently attached to cysteine residues via thioester linkages(*25–28*). There is growing evidence that interactions between FASN and palmitoylation targets are necessary for this protein modification(*28–30*). Thus, our discovery of a FASN-GSDMD interaction prompted us to examine whether GSDMD can be palmitoylated. We transiently transfected HEK293T cells with a full-length GSDMD construct and assessed palmitoylated GSDMD levels after 24 h using an acyl-biotin exchange (ABE) assay (**fig. S1d**). Indeed, recombinant GSDMD was robustly palmitoylated in HEK293T cells, and pretreatment with hydroxylamine (HAM), which cleaves the thioester bond to generate a sulfhydryl group, was essential for detecting signal (**fig. S1e)**. Silencing FASN with a specific *FASN* siRNA but not control scrambled siRNA significantly reduced GSDMD palmitoylation (**fig. S1f)**. Collectively, our results reveal an interaction between GSDMD and FASN and demonstrate that FASN is required for GSDMD palmitoylation.

### LPS triggers both GSDMD-FL and GSDMD-NT palmitoylation in macrophages

To identify which GSDMD region is palmitoylated, we transiently transfected HEK293T cells with GSDMD-FL, GSDMD-NT, and GSDMD-C terminal fragment (GSDMD-CT). Western blot analysis confirmed that both GSDMD-FL and GSDMD-NT but not GSDMD-CT were palmitoylated in 293T cells (**Fig. 1a**). Signals were significantly attenuated when cells were pretreated with the palmitate analog 2-bromopalmitate (2BP), an electrophilic α-brominated fatty acid widely used to inhibit protein palmitoylation, confirming signal derivation from protein palmitoylation (**Fig. 1a**). To examine palmitoylation of endogenous GSDMD, PMA-differentiated macrophage-like THP-1 cells were treated with or without LPS for 3 h and then nigericin for 35 min (LPS/Nig) to activate inflammasomes. Proteins were collected from the cell lysate and supernatant and palmitoylation was detected using the ABE method. Before LPS stimulation, GSDMD palmitoylation was virtually undetectable, and GSDMD palmitoylation was only detected in LPS/Nig-treated THP-1 cells. Similar to in HEK293T cells, both full length and cleaved N-terminal GSDMD were palmitoylated, and 2BP inhibited the palmitoylation (**Fig. 1b-c**). Similar results were also observed in mouse bone marrow-derived macrophages (mBMDMs), with palmitoylation of full length and cleaved N-terminal GSDMD only detected upon LPS/Nig stimulation (**Fig. 1d**). Of note, palmitoylation could be triggered by LPS alone, independent of Nig-induced inflammasome activation (**fig. S2**), so GSDMD palmitoylation in macrophages is not constitutive and relies on specific stimuli, consistent with a previous study showing that LPS upregulates palmitoyl enzyme transferases in macrophages(*31*). LPS-induced ROS production appeared to be critical for GSDMD palmitoylation although ROS alone could not trigger such an event (**Fig. 1c**). NF-kB activation was also involved since a specific NF-kB inhibitor significantly suppressed LPS-induced GSDMD palmitoylation (**Fig. 1c**). Our data also suggest that GSDMD cleavage is independent of its palmitoylation and 2BP-mediated inhibition of GSDMD palmitoylation does not affect cleavage (**Fig. 1b-c**), indicating that palmitoylation may be important for processes other than GSDMD cleavage.

**Figure 1.**
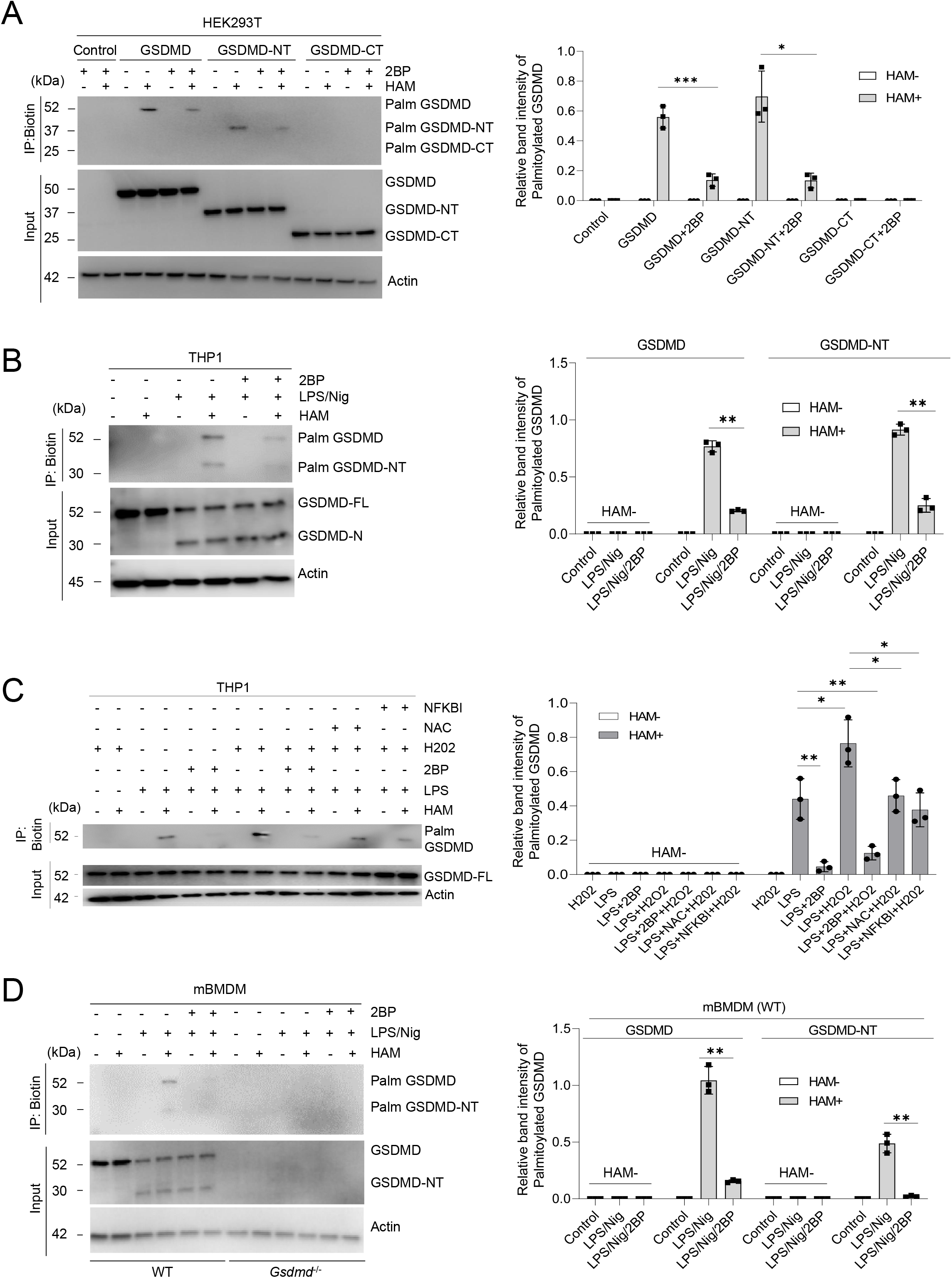
Both full length (GSDMD-FL) and cleaved GSDMD-NT are palmitoylated. **a,** HEK293T cells were transfected with or without GSDMD-FL, GSDMD-N, or GSDMD-C in the presence or absence of 2BP (50 μM). Cell lysates were treated with or without hydroxylamine, and immunoblot analyses were performed with anti-FLAG antibodies. Right, image represents the quantification of palmitoylated GSDMD proteins normalized to input. **b,** PMA-differentiated macrophage-like THP-1 cells were treated with or without LPS for 3 h, followed by 2-bromopalmitate treatment for 30 mins. Nigericin was then added to the cells and cell lysates subjected to the ABE assay in the presence or absence of hydroxylamine. Western blot analyses were performed with anti-GSDMD antibodies. The image to the right represents the quantification of palmitoylated GSDMD normalized to input. **C,** PMA-differentiated macrophage-like THP-1 cells were treated with or without LPS for 3 h, followed by pre-treatment with H202 for 30 min. 2BP (10 μM), N-acetyl cysteine (3 mM), JSH-23 (50 μM) was then added to the cells after H202 treatment for 30 min and cell lysates subjected to the ABE assay in the presence or absence of hydroxylamine. The image to the right represents the quantification of palmitoylated GSDMD normalized to input. **d,** Primary mouse bone marrow-derived macrophages were cultured in the presence of LPS (1 μg/ml) (3 h), followed by 2BP 10 μM (30 min) and treatment with nigericin (20 μM) for 1 h. Cell lysates were collected and treated with or without HAM and subjected to the ABE-palmitoylation assay. Right, palmitoylated GSDMD proteins were quantified by normalization to input. Data are mean ± SEM (A-C). Western blots shown are representative of three independent experiments. **P<0.01, ***P<0.001, unpaired Student’s *t*-test.

### GSDMD palmitoylation is critical for macrophage pyroptosis

To determine whether protein palmitoylation modulates pyroptosis, we treated PMA-differentiated macrophage-like THP-1 cells with two known palmitoylation inhibitors, cerulenin and *2BP(32-34*). LPS/Nig-induced cell death was assessed using SYTOX Green, a high-affinity nucleic acid dye that only penetrates dying cells with compromised plasma membrane integrity. Inhibition of palmitoylation with cerulenin or 2BP significantly suppressed pyroptosis as measured by SYTOX (**Fig. 2a-b**), the LDH cytotoxicity assay (**Fig. 2c**), or a luminescence-based cell viability assay to quantify the reducing potential as a surrogate of metabolically active cells (**Fig. 2d**). Furthermore, the two palmitoylation inhibitors significantly suppressed cell death-associated IL-1β release (**Fig. 2e**). The depalmitoylation inhibitor palmostatin B (PMB) did not affect cell death (**Fig. 2a-e**), indicating that depalmitoylation might not be a prominent regulator of macrophage pyroptosis. Cerulenin can also alter expression of NLRP3, IL-1β, and FASN in macrophages(*35*). We also noticed that cerulenin significantly suppressed caspase-1 activity in THP1 cells, further indicating that its effect on macrophage pyroptosis may not be a direct consequence of GSDMD palmitoylation (**Fig. 2f**). Thus, we included 2BP in all subsequent experiments as a more specific inhibitor of palmitoylation not affecting caspase-1 activity (**Fig. 2f**) and GSDMD cleavage (**fig. S3a-b**) but inhibiting LPS/Nig-induced cell death in a dose-dependent manner (**Fig. 2g-h**).

**Figure 2.**
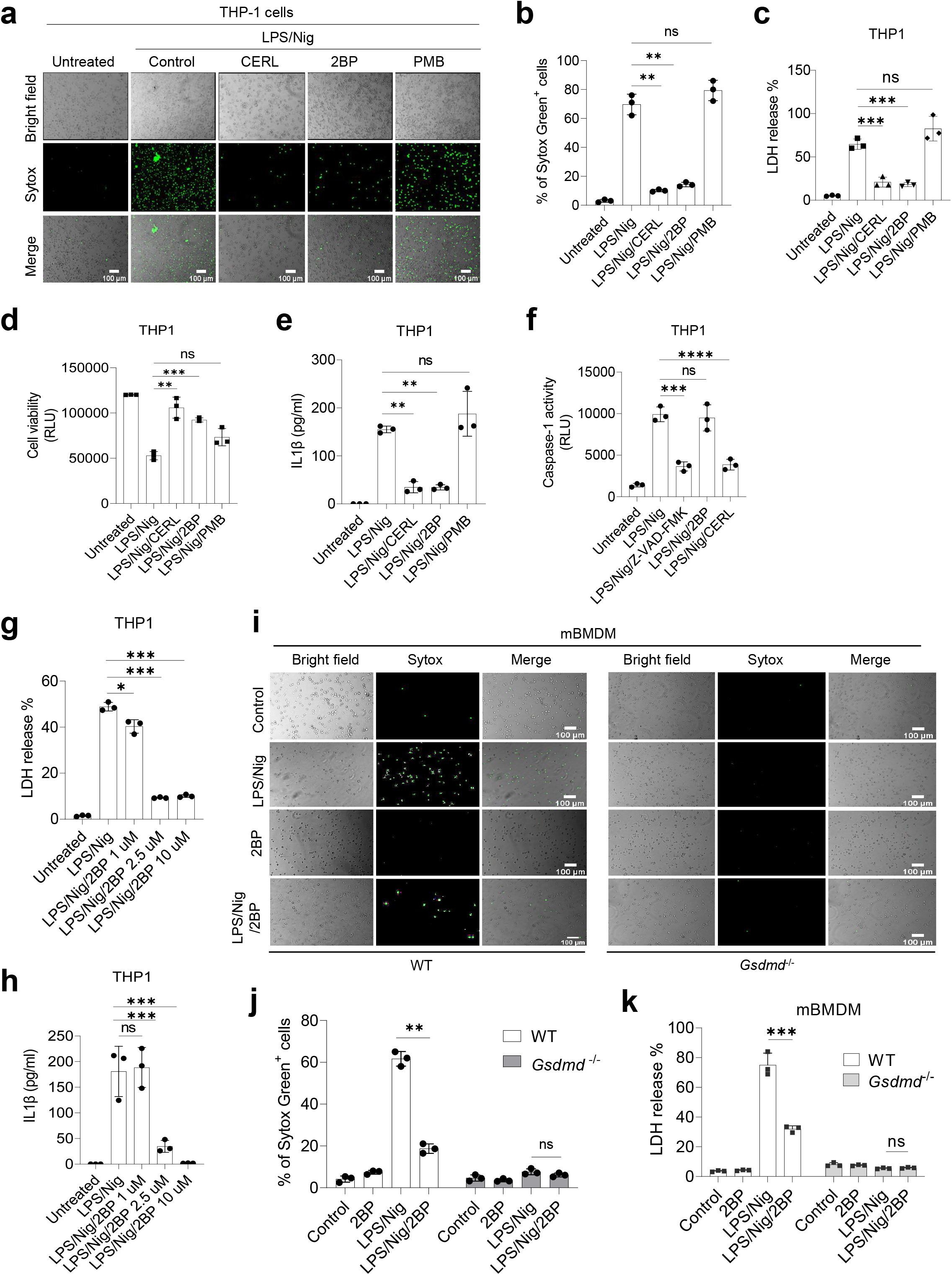
Inhibition of palmitoylation impairs pyroptosis in macrophages. **a,** Representative fluorescence images of THP-1 cells primed with LPS (3 h) followed by treatment with palmitoylation inhibitors (cerulenin, 50 μM; 2-bromopalmitate, 10 μM) and de-palmitoylation inhibitor (palmostatin B, 50 μM) for 30 min before stimulation with nigericin for 35 min. To avoid impacting transcription, inhibitors were added after LPS treatment. Pyroptotic pore formation was assessed with the SYTOX Green dye binding assay using ImageJ software. Scale bars, 100 μM. **b,** Quantification of relative SYTOX uptake in THP-1 cells. **c,** Cell culture media supernatants were diluted at least 100 times and LDH release was measured with a luminometer. **d,** Relative cell viability was calculated using the RealTime-Glo MT Cell Viability Assay (Promega). **e,** IL-1β secretion by THP-1 cells was measured by ELISA. **f,** The relative abundance of active caspase in cell culture supernatants of THP-1 cells was quantified with Caspase-1 Glo reagent (Promega). Luminescence was read with a luminometer. **g,** THP-1 cells treated with or without LPS/Nig in the presence or absence of 2BP were treated at different concentrations (1, 2.5, 10 μM). The resulting supernatants were quantified for LDH release. **h,** The level of IL-1β release from THP-1 cells was detected using an ELISA kit. **i,** Primary mouse bone marrow-derived macrophages (mBMDM) from wild type and *Gsdmd^-/-^* mice were primed with or without LPS for 3 h in the presence or absence of 2BP for 30 min before stimulation with nigericin for 1 h. Pyroptotic pore formation was assessed with the SYTOX Green assay, and fluorescence images were captured and quantified using ImageJ. Scale bars, 100 μM. **j,** Quantification of relative SYTOX uptake in mBMDMs. **k,** LDH release from WT and *Gsdmd^-/-^* cell culture supernatants was assessed using an LDH assay kit. **(a-k)** All data are representative of three independent experiments. Data are mean ± SEM. *P<0.05, **P<0.01, ***P<0.001, unpaired Student’s *t*-test.

Both SYTOX Green uptake (**Fig. 2i-j**) and LDH release (**Fig. 2k**) assays confirmed that inhibition of protein palmitoylation by 2BP also suppressed LPS/Nig-induced cell death in primary mBMDMs. GSDMD is crucial for pyroptosis, and LPS/Nig-induced pyroptosis was drastically reduced in GSDMD-deficient mBMDMs. 2BP did not further inhibit the death of GSDMD-deficient mBMDMs, suggesting that its effect is specific for GSDMD-mediated cell death (**Fig. 2i-k**).

Both full length GSDMD and the cleaved N-terminal GSDMD fragment can be palmitoylated. We next investigated whether the effect of GSDMD palmitoylation on cell death is directly mediated by the cytotoxic, pore-forming activity of GSDMD-NT. HEK293T cells expressing GSDMD-NT were treated with 2BP, and GSDMD-NT-induced cell death was measured by propidium iodide (PI) uptake and LDH release. 2BP treatment significantly reduced GSDMD-NT-induced lytic cell death in both assays (**fig. S4a-b**) without affecting GSDMD-NT expression (**fig. S4c**). We also established an inducible system in which recombinant human GSDMD-NT protein was expressed in a doxycycline (Dox)-inducible manner in HAP1 cells constitutively expressing EGFP. The HAP1 cell line is derived from human chronic myelogenous leukemia (CML) KBM-7 cells(*36*) with a near-haploid karyotype except for chromosomes 8 and 15. HAP1 cells are more sensitive to Dox induction than HEK293 and THP1 cells. GSDMD-NT-mediated cell death was assessed by measuring GFP fluorescence intensity, PI staining (**fig. S5a-b**), LDH release (**fig. S5c**), and cellular reducing potential (**fig. S5d**). Dox-induced GSDMD-NT expression induced prominent cell death according to all four metrics. Treatment with palmitoylation inhibitors rescued cells from pyroptotic cell death. In this inducible system, palmitoylation inhibitors did not alter GSDMD-NT expression, with similar amounts of recombinant protein expressed in untreated and inhibited cells (**fig. S5e**). Palmitoylation inhibition also did not affect Dox-induced GFP expression, again indicating that palmitoylation does not alter transcription or translation (**fig. S5f**). Our data suggest that palmitoylation is required for GSDMD-mediated pyroptosis, likely by facilitating GSDMD-NT activation.

### GSDMD palmitoylation occurs on the critical Cys192 residue and dictates GSDMD-NT membrane translocation

Palmitoylation often occurs on cysteine residues and less frequently on serine and threonine residues. To map the palmitoylation site in GSDMD-NT, we mutated each cysteine in GSDMD-NT and assessed palmitoylation using the ABE method (**Fig. 3a**). GSDMD-NT palmitoylation only significantly decreased when Cys192 but not other cysteine residues was mutated (**Fig. 3b**). We also treated cells with an alkyne-tagged palmitic acid analogue and subsequently labeled palmitoylated protein via a click chemistry conjugate(*37*) (**fig. S6a**). Palmitoylated GSDMD-NT levels were higher in wild type and Cys39, Cys57, Cys77, Cys122, and Cys265 mutants, but the palmitoylation signal was much weaker in the Cys192 mutant (**fig. S6b**). As expected, HAM could cleave the thioester bond and remove the palmitoyl moiety from the protein. These data highlight that Cys192 is crucial for palmitoylation.

**Figure 3.**
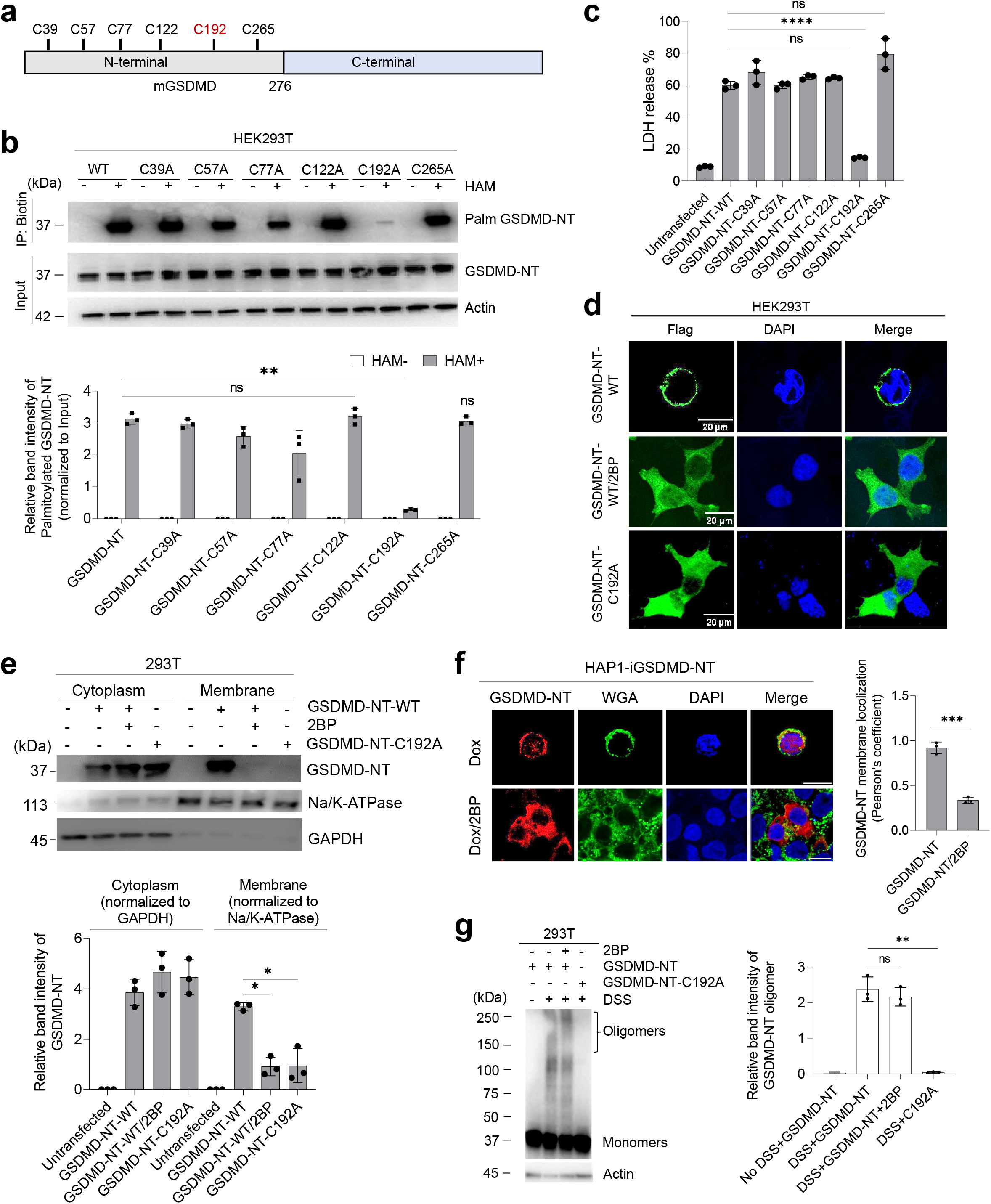
GSDMD is palmitoylated at Cys192 and its mutation abolishes GSDMD plasma membrane localization and pyroptosis. **a,** A schematic of the number of cysteine residues present in mouse GSDMD-NT. Highlighted in red is a potential cysteine site in GSDMD-NT that is palmitoylated. **b,** 293T cells were transfected with wild type GSDMD-NT or mutant N-GSDMD (Cys39A, Cys57A, Cys77A, Cys122A, Cys192A, Cys265A). After 24 h of transfection, cell lysates and supernatant were collected and pre-blocked with 20 mM NEM. Then, protein samples were precipitated with methanol:chloroform followed by treatment with 0.5 M hydroxylamine and biotin-azide. Streptavidin-pulled down samples were eluted with 2x Laemmli buffer containing reducing agent. Western blot analyses were performed with anti-FLAG HRP antibody. Palmitoylated proteins were quantified and normalized to corresponding input. **c,** LDH release from the cell culture media of wild type and mutant GSDMD-NT were quantified using an LDH assay kit (Promega). The luminescence signal was captured with a luminometer. **d,** Subcellular localization of GSDMD-NT in 293T cells treated with or without 2BP and mutant was analyzed by confocal microscopy. Scale bars, 20 μM. Images shown are representative of three independent experiments. **e,** Distribution of GSDMD-NT and mutant in subcellular fractions of 293T cells subjected to immunoblot analysis. Quantification of relative GSDMD-NT levels in these fractions. **f,** Subcellular localization of GSDMD-NT in Dox-induced GSDMD-NT cells were treated with or without 2BP and images were taken by confocal microscopy. Scale bars, 20μM. Images shown are representative of three independent experiments. **g,** Inhibition of palmitoylation did not affect GSDMD oligomerization. Oligomerization with transient expression of GSDMD-NT and mutant after treatment with or without 2BP (50 μM) was assessed by western blot analysis with anti-FLAG antibodies. Right, image represents the relative quantification of GSDMD-NT oligomers in 293T cells. Western blot images are representative blots from three independent experiments. Data are mean ± SEM. **P<0.01, ***P<0.001, ****P<0.0001, unpaired Student’s t-test.

Cells transfected with mutant GSDMD-NT were examined for LDH release. Wild type (WT) GSDMD-NT induced LDH release, whereas Cys192A, but not other mutations, significantly decreased LDH release (**Fig. 3c**). Palmitate acts as a hydrophobic membrane anchor, stably attaching palmitoylated protein to the cellular membrane(*38, 39*). Thus, we examined whether GSDMD-NT palmitoylation is required for its membrane localization. HEK293T cells were transiently transfected with WT or Cys192A-mutant GSDMD-NT and then treated with or without 2BP. GSDMD-NT localization was determined by both immunofluorescence imaging (**Fig. 3d**) and subcellular fractionation (**Fig. 3e**). Consistent with previous studies(*10, 18*), GSDMD-NT was predominantly membrane localized whereas GSDMD-NT-Cys192A was cytosolic. However, WT GSDMD-NT relocated to the cytoplasm upon 2BP treatment (**Fig. 3d-e**), suggesting that palmitoylation is required for membrane localization of GSDMD-NT for pyroptosis induction. Similar results were observed when recombinant human GSDMD-NT protein was expressed in a Dox-inducible manner in HAP1 cells (**Fig. 3f**). Interestingly, although full length GSDMD could be palmitoylated, its location remained cytosolic in HEK293 cells (**fig. S7a**) and both unstimulated and LPS-stimulated THP-1 cells (**fig. S7b**). Thus, palmitoylation only dictated membrane translocation of cleaved GSDMD-NT.

Next, we investigated the effect of palmitoylation on GSDMD-NT oligomerization. Western blotting revealed that WT GSDMD-NT, but not the Cys192 mutant, formed oligomers. Intriguingly, 2BP did not significantly reduce oligomerization (**Fig. 3g**), indicating that GSDMD-NT membrane translocation and oligomerization were two separate processes and that oligomerization did not rely on membrane translocation. Collectively, these findings corroborate the importance of S-palmitoylation in mediating GSDMD-NT localization and pore formation in pyroptosis.

### ZDHHC 5/9 palmitoylates GSDMD

Palmitoylation is mediated by palmitoyl acyl transferase (PAT) enzymes, a family of zinc finger and DHHC motif-containing (ZDHHC) proteins that transfer palmitate onto the thiol group of cysteine from cytosolic palmitoyl-CoA. Acyl protein thioesterases (APTs; depalmitoylation enzymes) can reverse this process(*40*) (**fig. S8a**). Twenty-four human DHHC domain proteins have been identified to date: 17 are known to have PAT activity(*40*), while the activity of other members (1, 11, 13, 14, 16, 23, and 24) is unknown. ZDHHC proteins have distinct substrate specificities. To identify the specific palmitoyl acyl transferases responsible for GSDMD palmitoylation and pyroptosis, we first identified which ZDHHC proteins were expressed in HEK293T cells. *ZDHHC* 1, 8, 11, 15, 19, and 22 mRNAs were almost undetectable by RT-qPCR in HEK293T cells (**fig. S8b**). We then co-transfected HEK293T cells with GSDMD-NT and siRNAs targeting each ZDHHC with PAT activity expressed in these cells, screening for siRNAs that suppressed GSDMD-NT-induced pyroptotic death as measured by an LDH release assay. ZDHHC5 and 9 were the most potent mediators of GSDMD palmitoylation (**fig. S8c**). We performed a similar screening in HAP1 cells expressing recombinant human GSDMD-NT protein in a Dox-inducible manner. Again, siRNAs targeting *ZDHHC5* and *9* most potently decreased LDH release compared with scrambled control siRNA (**fig. S8d**). The ABE assay revealed that silencing *ZDHHC5* or/and *9* significantly reduced GSDMD-NT palmitoylation compared with cells transfected with control scrambled siRNA (**fig. S8e-g**). Consistently, both ZDHHC5 and 9 were required for maximum LDH release, further demonstrating that GSDMD needs both palmitoyl enzymes to induce pyroptosis (**fig. S8h**). Additionally, fluorescent imaging showed that GSDMD-NT-induced PI uptake was inhibited when either *ZDHHC5* or 9 was knocked down (**Fig. S8i**). Of note, knocking down another ZDHHC protein, ZDHHC2, only moderately reduced GSDMD-NT-induced cell death in the LDH assay (**fig. S8c-d**) but had no detectable effect on PI uptake (**fig. S8i**). To further investigate the regulation of GSDMD palmitoylation by ZDHHC5 and 9, we overexpressed the enzymes in HEK293T cells and measured GSDMD palmitoylation with the ABE assay. Compared with control samples, GSDMD palmitoylation significantly increased in cells co-expressing ZDHHC5 (**fig. S9a**) or ZDHHC9 (**fig. S9b**).

### ZDHHC5/9-mediated GSDMD palmitoylation is critical for macrophage pyroptosis

To validate the physiological significance of ZDHHC5/9-mediated GSDMD palmitoylation, we first examined PMA-differentiated macrophage-like THP1 cells. THP1 cells expressed both ZDHHC5 and ZDHHC9, and LPS stimulation slightly upregulated their expression (**fig. S10a**). Upon stimulation with LPS/Nig, both full length and cleaved GSDMD-NT were palmitoylated, and siRNA knockdown of *ZDHHC5* or/and *9* significantly suppressed this palmitoylation (**Fig. 4a-c**). As a result, cells transfected with *ZDHHC5* or *9* siRNA released less LDH (**Fig. 4d**) and retained less SYTOX Green nucleic acid stain (**Fig. 4e**) upon LPS/Nig stimulation. Silencing both *ZDHHC5* and *9* led to almost complete block of GSDMD-NT palmitoylation and more significant reductions in LPS/Nig-induced pyroptotic death compared with silencing each ZDHHC individually (**Fig. 4c-e**). We also tried to knock down *Zdhhc5* and *9* with specific siRNAs in mouse primary BMDMs. Interestingly, SYTOX Green nuclear staining and LDH release were not reduced by *Zdhhc9* siRNA in mBMDMs **(fig. S10b-c**), whereas silencing *Zdhhc5* significantly suppressed pyroptotic cell death. ZDHHC9 expression was barely detectable in mBMDMs, potentially explaining why ZDHHC9 knockdown was ineffective (**fig. S10d**). Collectively, ZDHHC5/9 are the major palmitoyl acyl transferases responsible for GSDMD palmitoylation in macrophages and represent a potential target for regulating pyroptosis in infection and inflammatory diseases.

**Figure 4.**
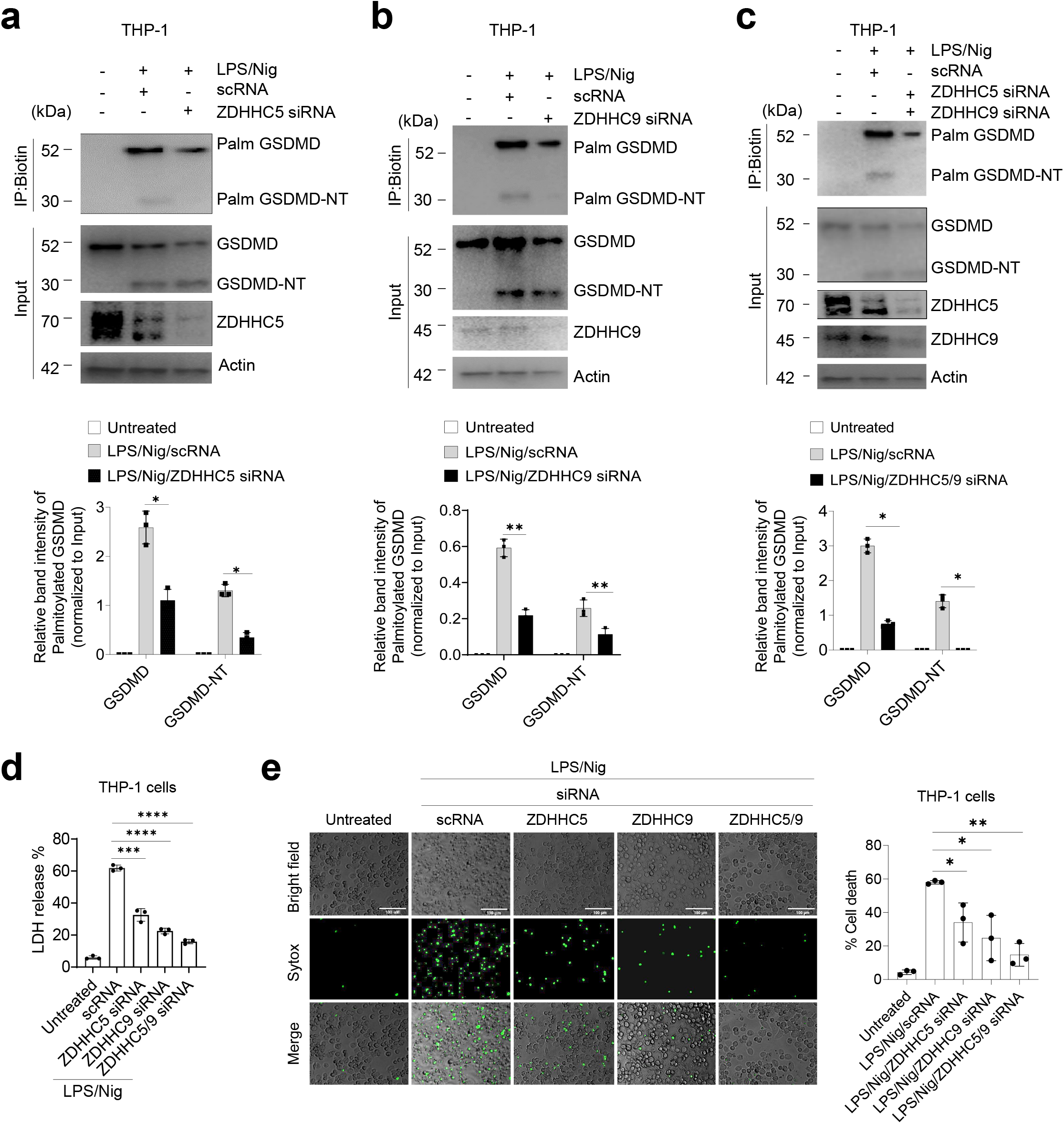
ZDHHC5/9-mediated palmitoylation is required for LPS/Nig-triggered pyroptosis in macrophages. **a-c,** THP-1 cells were transiently transfected (using a Lonza nucleofection kit) with control or siRNA targeting (a) *ZDHHC5*, (b) *ZDHHC9*, and (c) *ZDHHC5/9* followed by priming with LPS for 3 h and treatment with nigericin for 35 min. The ABE assay and immunoblot analysis were then performed with anti-GSDMD antibodies. Bar graph depicts the quantification of palmitoylated GSDMD protein normalized to relative input. n=3 mice per group. **d,** LDH release was measured from the cell culture supernatant of THP-1 cells. **e,** Fluorescence images are representative images of THP-1 cells expressing control and *ZDHHC* siRNA. The relative fluorescence signal of SYTOX Green uptake was captured by fluorescence microscopy. Quantification of relative SYTOX uptake in THP-1 cells. Data are representative of three independent experiments. *P<0.05, **P<0.01, ***P<0.001, unpaired Student’s *t*-test.

### Inhibition of palmitoylation extends the survival of mice with sepsis

GSDMD is an important innate immunity mediator, and GSDMD-deficient mice are protected from sepsis(*3, 18, 41*). To elucidate the potential role of GSDMD palmitoylation in sepsis, C57BL/6J mice were intra-peritoneally injected with LPS (25 mg/kg) and observed continuously for 7-10 days. For 2BP treatment, mice were first injected with 2BP (30 mg/mg) followed by LPS. To confirm GSDMD palmitoylation *in vivo* in septic mice, peritoneal cells from LPS-challenged control and 2BP-pretreated mice were collected after 12 h, and GSDMD palmitoylation was assessed using the ABE assay. GSDMD was robustly palmitoylated upon LPS stimulation, whereas 2BP treatment significantly reduced palmitoylation (**fig. S11a**). To assess the severity of sepsis, the overall inflammatory response was evaluated by assessing the concentrations of both serum and peritoneal fluid inflammatory cytokines by enzyme-linked immunosorbent assay (ELISA). Levels of tumor necrosis factor-α (TNF-α), interleukin-6 (IL-6), IL-1β, and IL-17 were drastically reduced in 2BP-treated mice (**fig. S11b**), as was LDH release (**fig. S11c**). Importantly, GSDMD disruption prevented all these phenomena, suggesting a specific role for GSDMD in mediating 2BP effects. Severe sepsis kills experimental mice. We therefore investigated whether 2BP treatment increased the survival rate of challenged mice. When challenged with 25 mg/kg LPS, all septic WT mice mock treated with phosphate buffered saline (PBS) died within two days. GSDMD-deficient mice and WT mice pretreated with 2BP had significantly better survival (>70%) than untreated WT mice (**fig. S11d**). When challenged with 54 mg/kg LPS, even GSDMD-deficient mice had a significant death rate, with about 40% of mice dying within three days. Pretreatment with 2BP failed to improve the survival of GSDMD-deficient mice (**fig. S11e**), suggesting that the 2BP-induced protective effects were largely mediated by GSDMD palmitoylation.

In addition to the LPS model, we also explored the effect of 2BP on sepsis induced by cecal ligation and puncture (CLP). Proinflammatory cytokine levels were low in unchallenged mice. After CLP, there was a significant increase in cytokine concentrations in wild-type peritoneal exudate, which was reduced by 2BP treatment (**fig. S11f**). In contrast, CLP-induced cytokine production was unaffected by 2BP treatment in *Gsdmd* KO mice (**fig. S11f**). To evaluate the extent of multi-organ dysfunction in these mice, circulating alanine transaminase (ALT), aspartate aminotransferase (AST), blood urea nitrogen (BUN), and creatinine levels were evaluated, which were all lower in 2BP-treated mice (**fig. S11g**). Similarly, peritoneal fluid LDH was reduced in 2BP-treated mice (**fig. S11h**). Furthermore, semi-quantitative histopathological examination of CLP-challenged mouse lung, kidney, and liver tissue revealed less tissue damage than mock-treated controls (**fig. S11i-j**). Consistent with the reduced organ damage, 2BP-treated mice showed significantly improved survival after CLP challenge, >55% survival compared with 0% in the mock-treated mice (**fig. S11k**). Similar to observations in the LPS sepsis model, all 2BP-induced protective effects were abolished in GSDMD-deficient mice. Taken together, these findings suggest that GSDMD palmitoylation is a key event regulating GSDMD and pyroptosis, highlighting the potential for targeting this mechanism to treat infectious and inflammatory diseases.

### A cell-permeable, GSDMD palmitoylation-specific competitive peptide suppresses macrophage pyroptosis and alleviates sepsis

As a general inhibitor of palmitoylation, 2BP targets multiple proteins. To specifically block GSDMD palmitoylation, we used a competitive peptide corresponding to the conserved GSDMD palmitoylation region (**Fig. 5a**), an approach commonly applied for target-specific palmitoylation *inhibition*(*42–46*). A cell permeable peptide (CPP), CPP-R9 (polyarginine), was fused to the GSDMD peptide (CPP-W) to achieve delivery to the cytosol of target cells. A peptide carrying a C192A substitution (CPP-M) was synthesized and used as a negative control (**Fig. 5b**). CPP-W, but not CPP-M, efficiently suppressed GSDMD palmitoylation (**Fig. 5c**) and consequently inhibited the death of Dox-inducible HAP1 cells expressing GSDMD-NT, as measured by GFP expression and PI staining (**Fig. 5d)**, LDH release (**Fig. 5e**), and cellular reducing potential (**Fig. 5f**). Similar pyroptosis-blocking effects were detected in mBMDMs challenged with LPS/Nig (**Fig. 5g-J**).

**Figure 5.**
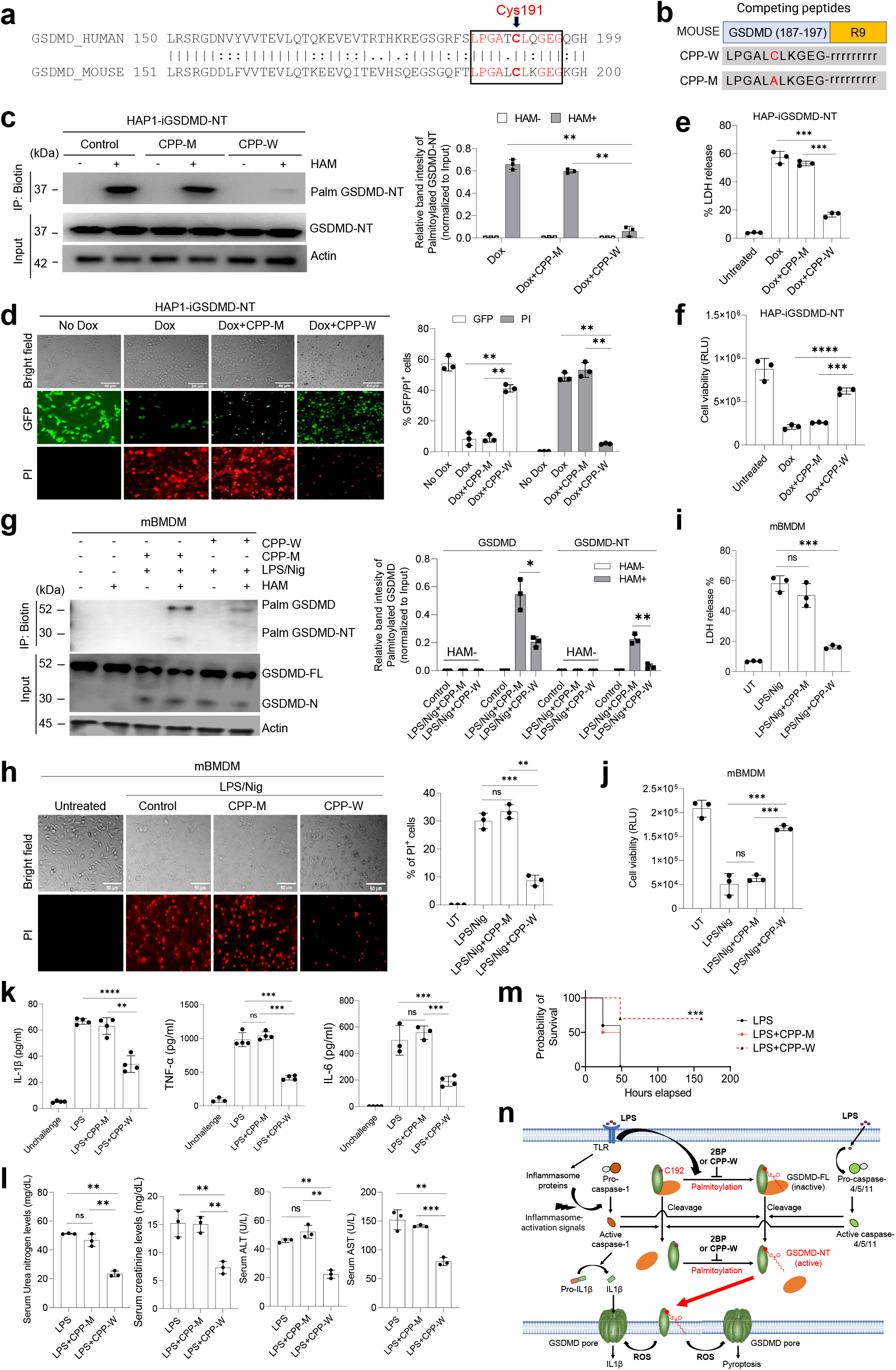
A cell permeable peptide specifically inhibits GSDMD palmitoylation, suppresses macrophage pyroptosis, and alleviates the severity of sepsis. **a,** Protein sequence alignment of human and mouse GSDMD using EMBOSS Matcher. The conserved cysteine residues are marked bold. **b,** Cell permeable, GSDMD-competing peptides. R9 represents poly-arginine. Lower case amino acid symbol (r) represents the D-form of arginine. **c,** Doxycycline-inducible HAP1 cells expressing GSDMD-NT (HAP1-iGSDMD-NT) were pretreated with CPP-M (50 μM) or CPP-W (50 μM) for 4 h followed by induction with doxycycline (100 ng/ml) for 3 h. GSDMD palmitoylation was measured using the ABE assay as described in **Fig. 2b**. Western blots shown are representative of three independent experiments. Right, palmitoylated GSDMD proteins were quantified by normalization to input. Data are mean ± SEM. *P<0.05, **P<0.01, ***P<0.001, unpaired Student’s *t*-test. **d,** Representative fluorescence images of GFP expression and PI uptake in i-GSDMD-NT cells treated with or without indicated peptides. Right, the relative mean fluorescence intensity of PI and GFP were quantified using ImageJ. Data are mean ± SEM (n=3). *P<0.05, **P<0.01, ***P<0.001, unpaired Student’s *t*-test. **e,** Cell culture media supernatants were subjected to the LDH assay and LDH percentage release was calculated for each group. **f,** The cells were subjected to a cell viability assay using the RealTime-Glo MT Cell Viability Assay kit (Promega). Relative luminescence units (RLU) were captured with a luminometer. **g,** Primary mBMDM cells were cultured in the presence of LPS (1 μg/ml) (3 h), followed by CPP-M (50 μM) or CPP-W (50 μM) for 1 h and treatment with nigericin (20 μM) for 1 h. Cell lysates were collected and treated with or without HAM and subjected to the ABE-palmitoylation assay. Western blots shown are representative of three independent experiments. Right, palmitoylated GSDMD proteins were quantified by normalization to input. Data are mean ± SEM (n=3). **P<0.01, unpaired Student’s *t*-test. **h,** Representative fluorescence images of PI uptake in mBMDM cells treated with or without indicated peptides. Right, the percentage of PI was quantified using ImageJ. Data are mean ± SEM (n=3). *P<0.05, **P<0.01, ***P<0.001, unpaired Student’s *t*-test. **i,** LDH release from cell culture supernatants was assessed using an LDH assay kit. Data are mean ± SEM (n=3). *P<0.05, **P<0.01, ***P<0.001, unpaired Student’s *t*-test. **j,** Cell survival was assessed using RealTime-Glo MT Cell Viability Assay kit (Promega). Data are mean ± SEM (n=3). *P<0.05, **P<0.01, ***P<0.001, unpaired Student’s *t*-test. **k,** C57BL/6J mice were pretreated with CPP-W (50 mg/kg, i.p.) or CPP-M (50 mg/kg, i.p.) for 6 h, followed by intraperitoneal challenge with LPS (20 mg/kg) for 12 h. Peritoneal fluid IL1-β,TNF-α, and IL6 levels were measured by ELISA (n=3-4 mice per group). *P<0.05, **P<0.01, ***P<0.001, unpaired Student’s I-test. **l,** Circulating AST, ALT, BUN, and creatine levels in the serum. n=3 mice per group. *P<0.05, **P<0.01, ***P<0.001, unpaired Student’s *t*-test. **m,** C57BL/6J mice were pretreated with CPP-W (50 mg/kg, i.p.) or CPP-M (50 mg/kg, i.p.) for 6 h, followed by intraperitoneal challenge with LPS (25 mg/kg). Survival rates were calculated using Kaplan-Meier survival curves. n=10 mice per group. *P<0.05, **P<0.01, ***P<0.001, log-rank test. **n,** Graphical illustration of GSDMD palmitoylation and its role in macrophage pyroptosis.

Finally, we investigated whether this GSDMD palmitoylation-targeting peptide alleviated the severity of LPS-induced sepsis *in vivo*. Indeed, due to a reduction in GSDMD-mediated macrophage pyroptosis, serum collected from CPP-W-treated septic mice contained lower levels of pro-inflammatory cytokines TNF-α, IL-1β, and IL-6, confirming attenuated systemic inflammation in these mice compared with PBS or CPP-M controls (**Fig. 5k**). CPP-W treatment also mitigated the extent of multi-organ dysfunction in septic mice, as evidenced by lower serum ALT, AST, BUN, and creatinine levels (**Fig. 5l**). Consistently, treatment with CPP-W significantly improved the survival of LPS-challenged mice. At a dose of 25 mg/kg LPS, all control and CPP-M-treated mice died within 48 h, whereas >70% CPP-W treated mice survived (**Fig. 5m**). Taken together, these findings suggest that specific inhibition of GSDMD palmitoylation could be an effective therapeutic strategy for sepsis.

## Discussion

Here we identified palmitoylation of GSDSD on Cys191/192 as a regulatory mechanism controlling GSDMD membrane localization and activation, providing a novel target for modulating immune activity in infectious and inflammatory diseases. In a mouse sepsis model, pharmacological inhibition of GSDMD palmitoylation with 2BP or a GSDMD-specific cell permeable competing peptide, CPP-W, protected mice from organ injury associated with exaggerated cytokine release. Mice with impaired GSDMD palmitoylation survived for longer with reduced circulating levels of pro-inflammatory cytokines IL-1β, TNF-α, and IL-6 than controls. The protective effect was largely mediated by GSDMD and was abolished in GSDMD-deficient mice.

GSDMD palmitoylation in macrophages was tightly regulated. Although GSDMD could be spontaneously palmitoylated in HEK293T and HAP1 cells, its palmitoylation in human macrophage-like THP1 cells and mouse bone marrow-derived macrophages only occurred after LPS stimulation. LPS-induced ROS were essential for GSDMD palmitoylation; GSDMD lipidation was significantly inhibited by scavenging ROS. Thus, GSDMD could be a key cellular sensor for ROS which have been implicated in many aspects of the immune response including regulation of inflammasome activity (*22, 47–49*) However, ROS appeared not to be sufficient to induce GSDMD palmitoylation and NF-kB-mediated cellular events were also required for LPS-elicited GSDMD lipidation.

How does palmitoylation dictate GSDMD function and regulate pyroptosis in macrophages? GSDMD-NT-mediated plasma membrane perforation triggers membrane rupture and subsequent lytic pyroptotic cell death, leading to release of pro-inflammatory cytokines IL-1β and IL-18(*1, 3, 4, 10*). GSDMD-NT forms pores at the plasma membrane through oligomerization. Both membrane translocation and oligomerization were thought to be important for GSDMD-NT-induced pyroptosis, although a GSDMD-NT Cys39 mutant appears to be defective in oligomerization but still elicits obvious cell death(*10, 50*). Cys191/192 is crucial for oligomerization(*10*). GSDMD-NT oligomerization can be inhibited by reducing agents, and Cys191/192 mutation impairs oligomerization. Accordingly, it has been suggested that intra- or intermolecular disulfide bonds between cysteine residues are critical for oligomerization. However, GSDMD crystal(*7*) and cryo-EM(*15*) structures have revealed that Cys191/192 is not adjacent to any Cys residues from neighboring GSDMD-NT subunits. Thus, the role of Cys191/192 in GSDMD oligomerization remains to be defined. Palmitoylation facilitated GSDMD-NT membrane translocation but did not appear to be required for oligomerization, which was not altered in cells treated with 2BP or CPP-W, indicating that GSDMD-NT membrane translocation and oligomerization are two separate processes. Indeed, it has also been shown that ROS facilitate GSDMD oligomerization but not membrane translocation(*22*). Membrane-associated GSDMD-NT monomers can be converted into pore-forming oligomers when exposed to ROS-inducing agents, suggesting that oligomerization can occur after membrane translocation. How the GSDMD-NT molecules are recruited to the membrane is poorly defined. Our results support a model in which palmitoylation of GSDMD mediates its membrane translocation (**Fig. 5n**). GSDMD-NT can be palmitoylated at Cys191/192, which targets it to plasma membrane. Disruption of palmitoylation at Cys191/192 blocks the membrane localization. Both immunofluorescence imaging and subcellular fractionation revealed that wild-type GSDMD-NT in CPP-W or 2BP-treated cells and GSDMD-NT harboring a Cys191/192 to Ala mutation failed to translocate to the plasma membrane. Therefore, although GSDMD-NT can directly bind to phosphatidylinositol phosphates and phosphatidylserine in the cell membrane inner leaflet, we argue that S-palmitoylation is also absolutely required to anchor cleaved GSDMD-NT to the membrane, facilitating plasma membrane rupture and pyroptotic cell death. Indeed, Cys191/192 in GSDMD is located in a disordered loop region and exposed to the solvent, making it easily accessible for palmitoylation(*7, 15*). Additionally, in the GSDMD-NT pore, Cys191/192 is right at the tips of the β-barrel that forms the pore, further supporting its involvement in anchoring the cleaved protein to the membrane(*15*) (**fig. S12**).

## Acknowledgements

The authors thank Tsan Sam Xiao, Jonathan Kagan, Marilyn D. Resh, Bryan C. Dickinson and Li Chai for helpful discussions and suggestions. For MS data collection and analysis, we thank Renne Robinson, Steven Kolakowski, Bogdan Budnik and Mei Chen at the HMS Proteomics Core Facility. This work was supported by National Institutes of Health grants 1R01AI142642, 1R01AI145274, 1R01AI141386, R01HL092020, and P01HL158688. A.H. was supported by NIH training grant T32HL066987.

## Author Contributions

H.R.L. and A.B. inputted conceptualization; H.R.L., A.B., A.H., F.L., and L.G. designed the methods; A.B., A.H., L.G., T.H., and F.L acquired samples; A.B., A.H., L.G., and T.H. performed analysis.; H.R.L., H.W., J.L., X.L., and R.X. provided resources; H.R.L. supervised all work. H.R.L. and A.B. wrote the paper with input from all authors. All coauthors read, reviewed, and approved the manuscript.

## Data availability

Mass spectrometric data have been deposited in ProteomeXchange Consortium with the dataset identifier PXD039133. Other datasets generated during and/or analyzed during the current study are available from the corresponding authors on reasonable request. Source data are provided with this paper.

## Lead contact

Further information and requests for resources and reagents should be directed to and will be fulfilled by the lead contact, Hongbo R. Luo.

