## Supplementary text and figures for "Palmitoylation of gasdermin D directs its membrane translocation and pore formation in pyroptosis"

### Supplementary Materials

This PDF file includes:

Materials and Methods
Supplementary Text
Figs. S1 to S12
Tables S1 and S2
References

#### Materials and Methods

##### Mice

Male C57BL/6J mice aged 6-8 weeks were purchased from the Jackson Laboratory (Bar Harbor, ME). *Gsdmd* KO mice(*1*) (on C57BL/6J background) were inbred and maintained in a pathogen-free system. For studies related to LPS or CLP-induced sepsis, 8–10-week-old male mice were used. For in vitro studies, mice of both sexes between 8 to 10 weeks old were used. All mice were housed and cared for in approved veterinary facilities located within Boston Children's Hospital, which provides sterile isolator cages with fresh food, water, and bedding weekly. All animal experiments were conducted in accordance with the Animal Welfare Guidelines of the Boston Children's Hospital. The Boston Children's Hospital Animal Care and Use Committee approved and monitored all procedures.

##### Cell culture

HEK293T and THP-1 cell lines were purchased from the ATCC. HAP1 was procured from Horizon Discovery (Cambridge, UK). HEK293T, THP-1, and HAP1 cells were maintained in DMEM, RPMI-1640, and IMDM media, respectively, containing 10% fetal bovine serum (FBS) and 1% penicillin/streptomycin.

##### LPS-induced sepsis model

Sepsis was induced in C57BL/6 mice (6-weeks old) and *Gsdmd* KO mice by intraperitoneal injection of LPS (25 mg/kg). Before LPS injection, mice were treated with 2-bromopalmitate (30 mg/kg). Blood and peritoneal cells were collected after 12 h. Serum was separated from blood by spinning down at 5,000 rpm for 5 min. Separated serum was diluted at least 1:5 and cytokines quantified by ELISA. Peritoneal cells were isolated and analyzed for LDH release, a palmitoylation assay, and inflammatory cytokine expression by ELISA.

##### Cecal ligation and puncture

Cecal ligation and puncture (CLP) was performed as previously(*2, 3*). Briefly, age- and sex-matched KO and WT mice were anesthetized with a mixture of xylazine (10 mg/kg) and ketamine (100-120 mg/kg). The surgical site was prepared by removing hair from the abdominal region using hair removal cream and sterilization using iodine and alcohol-soaked pads. A ventral midline incision was made on the skin and then on the peritoneum to expose the cecum, which was located using forceps and tied about 1-1.5 inches from its blunt end. Next, the cecum was perforated with an 18G needle twice to induce severe sepsis. The cecum was placed in its original position after squeezing a small amount of cecal contents into the peritoneal cavity. The peritoneum was sutured, and tissue clips were used to appose the skin. One ml of normal saline was injected into each mouse subcutaneously to compensate for fluid loss during surgery.

**Reagents and antibodies**

Protease inhibitor cocktail and PMSF were purchased from Thermo Fisher Scientific (Waltham, MA; 78440 and 36978). N-ethylmaleimide (04259), cerulenin (C2389), 2-bromopalmitate (21604), hydroxylamine solution (467804), and palmostatin B (178501) were from Sigma-Aldrich (St. Louis, MO). Biotin HPDP (16459), biotin azide (13040), and palmitic acid alkyne (13266) were purchased from Cayman Chemical Company (Bar Harbor, ME). The click chemistry protein reaction buffer kit (C10276) was from Thermo Fisher Scientific. Rabbit polyclonal gasdermin-D antibody (39754) was purchased from Cell Signaling Technology (Danvers, MA). Goat anti-rabbit IgG secondary antibody HRP was from Thermo Fisher Scientific (65-6120). Anti-FLAG M2 affinity gel (A2220), anti-β-actin HRP (A3854), and monoclonal anti-FLAG M2 HRP (A8592) were from Sigma-Aldrich. Streptavidin agarose (20353) was from Thermo Fisher Scientific. Ultrapure LPS and nigericin were purchased from InvivoGen (San Diego, CAA). HA-tag mouse mAb HRP antibody (2999S) and fatty acid synthase rabbit mAb (3180S) were from Cell Signaling Technology. ZDHHC5 polyclonal antibody (A304-652A), Lipofectamine 3000 (L3000008), ZDHHC9 polyclonal antibody (PA5-26721), goat anti-mouse IgG (H+L) secondary antibody Alexa Fluor 488 (A32723), goat anti-rabbit IgG secondary antibody Alexa Fluor 594 (A-11012), and wheat germ agglutinin Alexa Fluor 647 conjugate (W32466) were from Thermo Fisher Scientific. SuperSignal West Femto Maximum Sensitivity Substrate (34096) was from Thermo Fisher Scientific.

**Cell Fractionation**

Cell cytoplasmic and membrane fractions were prepared using a Cell Fractionation Kit (Cell Signaling Technology). Briefly, HEK293T cells (2.2 x $10^6$) were seeded in a 100-mm cell culture plate. After 24 h of transfection with mGSDMD-NT or mGSDMD-C192A cDNA, cells were washed with PBS, trypsinized, and then collected by centrifugation at 400g for 5 min. Cells were then lysed with 500 µl of cytoplasmic isolation buffer, vortexed, and incubated on ice for 5 min followed by centrifugation at 500g for 5 min. The supernatant was collected as a cytoplasmic fraction. The remaining pellet was resuspended in 500 µl of membrane isolation buffer and incubated on ice for 5 min. After centrifugation at 8,000g for 5 min, the supernatant was collected and stored as a membrane fraction.

**Cell-permeable peptide synthesis**

Cell-permeable GSDMD competing and control peptides corresponding to mouse GSDMD (aa 187-197) were chemically synthesized and conjugated with CPP-R9 (D-form) at LifeTein (Somerset, NJ). The purity and sequence of the peptides were verified by mass spectrometry.

**Cloning and mutagenesis**

cDNA clones from mouse and human *GSDMD* were cloned into pCDNA vector. FLAG-gasdermin D-C-terminus was purchased from Genscript Biotech Corp (Piscataway, NJ). HA-tagged DHHC 1-23 plasmids were obtained from M. Fukata, Japan. FLAG-N-*Gsdmd* wild type and FLAG-*Gsdmd*-NT mouse mutants C39A, C57A, C77A, C122A, C192A, and C265A were generated as previous described(*4*).

**Co-immunoprecipitation**

Cells were washed with PBS and lysed with lysis buffer (50 mM Tris pH 7.4, 150 mM NaCl, 1 mM EDTA, 1% NP-40) containing protease inhibitor and incubated for 30 min in ice. Briefly, cell lysates were centrifuged at 16,000 rpm for 10 min. Protein in the supernatant was quantified with a Bradford assay. Cell lysate with or without anti-FLAG M2 agarose beads were incubated overnight at 4°C with gentle rocking. After 24 h, samples were spun down and washed at least three times with wash buffer. Proteins were eluted with 0.2 M glycine (pH 2.0) and samples were processed for mass spectrometry and western blotting.

**Mass spectrometry**

Immunoprecipitated protein samples were concentrated using a speed vacuum concentrator and samples were dissolved in 100 µl TEAB (triethylammonium bicarbonate), heated to 95°C for 5 min, and cooled to room temperature (RT). In-solution proteins were digested with trypsin/LysC (1:50 ratio) and incubated at 37°C for 3 h. Digested proteins were desalted using a c18 spin column. Samples were injected into an Orbitrap Lumos mass spectrometer equipped with a dual pump Ultimate Nano LC (Thermo Fisher, San

Jose, CA). Peptides were separated onto a 100 μm inner diameter microcapillary trapping column packed first with approximately 5 cm of C18 Reprosil resin (5 μm, 100 Å, Dr. Maisch GmbH, Germany) followed by a 50 cm analytical uPAC column by PharmaFluidics (Ghent, Belgium). Separation was achieved by applying a gradient from 5-27% ACN in 0.1% formic acid over 90 min at 200 $nl.min^{-1}$. Electrospray ionization was enabled by applying 1.8 kV using a home-made electrode junction at the end of the microcapillary column and sprayed from fused silica pico tips (New Objective, Littleton, MA). The LTQ Orbitrap Lumos was operated in data-dependent mode for mass spectrometry. The mass spectrometry survey scan was performed in the range of 395-1,800 m/z at a resolution of 6 × 104, followed by selection of the thirty most intense ions (TOP30) for CID-MS2 fragmentation in the ion trap using a precursor isolation width window of 2 m/z, AGC setting of 10,000, and a maximum ion accumulation of 200 ms. Singly-charged ion species were not subjected to CID fragmentation. The normalized collision energy was set to 35 V and an activation time of 10 ms. Ions in a 10 ppm m/z window around ions selected for MS2 were excluded from further selection for fragmentation for 60 s.

**MS data analysis**

Raw data were submitted for analysis in Proteome Discoverer 2.4 (Thermo Fisher Scientific) software. MS/MS spectra were assigned using the SEQUEST HT algorithm by searching the data against a protein sequence database including all entries from our Uniprot_HUMAN_SPonly_2018.fasta database as well as other known contaminants such as human keratins and common lab contaminants. SEQUEST HT searches were performed using a 20-ppm precursor ion tolerance and requiring each peptide's N-/C-termini to adhere with trypsin protease specificity, while allowing up to two missed cleavages. An MS2 spectra assignment false discovery rate (FDR) of 1% at both the protein and peptide level was achieved by applying the target-decoy database search. Filtering was performed using Percolator (64-bit). Peptide N termini and lysine residues (+229.162932 Da) were set as static modifications, while methionine oxidation (+15.99492 Da) was set as a variable modification.

**Detection of GSDMD palmitoylation by acyl biotin exchange (ABE)**

To detect palmitoylation, cells were lysed with lysis buffer (50 mM Tris pH 7.2, 150 mM NaCl, 1% NP-40, 1 mM EDTA) with 20 mM N-ethylmaleimide and protease inhibitor. Samples were incubated for 2 h with agitation at 4°C and centrifuged at 16,000 x g for 15 min. The protein concentration was determined prior to ABE. 200 μL of protein samples were precipitated with methanol:choloroform:water, and the resulting pellet was dissolved in lysis buffer with or without 0.5 M hydroxylamine for 1 h at RT. After incubation, proteins were precipitated again and incubated with 4 mM biotin HPDP for 1 h. Precipitated proteins were dissolved in 1% SDS containing lysis buffer and diluted 1:10 with PBS, and 30 µl of streptavidin agarose beads were added and incubated overnight at 4°C. Beads were washed four times with lysis buffer and proteins were mixed with 2x SDS-Laemmli buffer containing 2-ME in a boiling water bath for 5 min. Proteins were separated by SDS-PAGE.

**Detection of GSDMD palmitoylation by click chemistry**

HEK293T cells were treated with alkyne-palmitic acid (20 µM) for 24 h in 10% charcoal-stripped FBS medium. Cell lysates were lysed in lysis buffer (50 mM Tris pH 7.2, 150 mM NaCl, 10% glycerol, 1% NP-40, 1 mM EDTA). Cell lysates were centrifuged at 16,000 x g for 5 min, the proteins were quantified, and samples were precipitated and subjected to a click chemistry reaction using the Click-iT Protein Buffer kit (Thermo Fisher Scientific, C10276) with 5 mM biotin azide and incubation for 30 min at RT. Then, proteins were precipitated again, and the pellets were incubated with or without hydroxylamine, and the samples were dissolved in 1% SDS containing lysis buffer and incubated with streptavidin agarose beads overnight. Proteins were eluted from beads with 2x Laemmli buffer containing reducing agents.

**siRNA knockdown of GSDMD**

THP-1 and primary bone marrow-derived macrophages (1 million cells/ml) were electroporated using Lonza Bioscience Nucleofector kit (Morrisville, NC; VCA-1003 and VPA-1009) with scramble control RNA and gene-targeting siRNAs for 48 h. For HEK293T cells, siRNA was transfected using Lipofectamine 3000.

**Western blotting**

Cell lysates were lysed in lysis buffer containing 25 mM tris pH 7.4, 150 mM NaCl, 1% NP-40, 10% glycerol, and 1mM EDTA. Samples were spun down at 16,000 x g for 10 min. Lysates were fractionated in 4-15% gradient SDS-PAGE (BioRad Laboratories, Hercules, CA), proteins were transferred to E-PVDF membranes (Millipore, Burlington, MA), and membranes were probed with primary antibodies (1:100) in 2% BSA in blocking buffer overnight at 4°C. After incubation, blots were washed three times with 1x TBST and probed with secondary HRP-conjugated antibody (1:10,000) for 1 h at RT. Membranes were washed with TBST and mixed with 1:1 SuperSignal West Femto Max substrate, and chemiluminescence signal was captured using a BioRad Chemidoc XP machine. Band intensities were quantified using ImageJ software. On each blot, the band with the strongest intensity was used for normalization to minimize inter-experimental variation (e.g. the signal intensity variation caused by different exposure time).

**LDH cytotoxicity assay**

Cell culture supernatants were centrifuged at 10,000 rpm for 5 min. Samples were diluted at least 1:100 times in LDH storage buffer. 100 µl of LDH substrate mix was added to the samples and incubated in the dark at RT for 30 min, and luminescence was captured using a Biotek luminometer. The percentage LDH release was calculated.

**Cytokine measurements**

WT and *Gsdmd* KO mice were intraperitoneally challenged with LPS (15 mg/kg) after 2-bromopalmitate treatment. Six hours later, retro-orbital blood was drawn and collected in Eppendorf tubes. The blood was centrifuged at 5000 rpm for 5 min to collect the serum. Peritoneal fluid was collected from the peritoneal cavity. IL-1β, TNF-α, IL-6, and IL-17A were measured using an ELISA Kit (Thermo Fisher Scientific).

**Immunofluorescence imaging**

To visualize the localization of GSDMD-NT, HEK293T cells were plated on coverslips and transiently transfected with wild-type FLAG *Gsdmd*-NT and mutant FLAG *Gsdmd*-N-C192A. After 6 h transfection, cells were treated with or without 2-bromopalmitate. 24 h later, cells were washed with PBS and fixed with 3% paraformaldehyde and stained with wheat germ agglutinin, followed by washing and permeabilization for 15 min with 0.5% triton. 2% BSA blocking buffer was added to the cells prior to incubation for 30 min. Goat anti-mouse FLAG primary antibody was diluted 1:2000 in blocking buffer and incubated at 4°C overnight. After a few washes, Alexa Fluor 488-conjugated secondary antibody (1:1000) was added and incubated further for 1 h at RT. Cells were washed and visualized using a Plan-Apochromat 63x/1.4 Oil M27 objective and processed with ImageJ. Colocalization efficiency was quantified with the ImageJ plugin coloc2 with background subtraction and ROI selection. Results are presented as normalized Pearson's correlation.

**Dox-inducible system (HAP1-iGSDMD-NT cells)**

cDNA encoding human gasdermin D N terminal region (1-275) was amplified and cloned into the pLVX Tet plasmid. To generate stable expression of GSDMD-NT, the pLVX tetone-hGsdmd-1-275-2A-GFP-IRES-Blast vector was transfected into HAP1 cells with Lipofectamine 2000 and selected with blasticidin for 24 and 48 h. Single cell clones were obtained and screened for GFP by FACS. To confirm gene expression of GSDMD-NT, cells were treated with doxycycline (10-100 ng/ml) for 1-6 h. After Dox stimulation, cell death was observed by fluorescence microscopy with PI staining. GFP-positive signal and PI-positive cells were captured and quantified using ImageJ. Media supernatants were collected and quantified for LDH release. HAP1 cells expressing inducible GFP (HAP1-iGFP) were used as a control for this study.

**Real-time PCR**

Total RNA was isolated using the Qiagen RNeasy Mini kit (Qiagen, Hilden, Germany). Using a cDNA synthesis kit (Bio-Rad), RNA was reverse transcribed into cDNA. cDNA was amplified using a Bio-Rad CFX 96 real-time PCR detection system. Gene expression was normalized to *Gapdh* loading control.

**Primary mouse bone marrow-derived macrophages and pyroptosis assay**

Wild type and *Gsdmd* knockout bone marrow was isolated as described previously. Bone marrow cells were isolated and seeded into six-well plates and maintained in DMEM containing 10% FBS, 100 U/ml penicillin/streptomycin, with 30 ng/ml recombinant M-CSF and incubated for 6 days. The medium was changed every three days. On day 6, adherent macrophages were washed once with DMEM, and cells were primed with LPS (1 μg/ml) for 3 h followed by secondary stimulation with nigericin 20 μm for 1 h. The supernatant was removed and quantified for LDH release. The remaining cells were stained with STOX green (1:30,000) and PI. Fluorescence signal was captured, and images were quantified using ImageJ.

**Histological staining and scoring**

Fresh lung, kidney, and spleen tissues were fixed in 4% paraformaldehyde (PFA) and then gradually dehydrated and embedded in paraffin. After that, 4 μm-thick tissue sections were cut and stained with hematoxylin and eosin (H&E) for further light microscopy observation. Lung injury scores were assessed based on the following parameters: alveolar capillary congestion, hemorrhage, edema, inflammatory cell infiltration, integrity, and alveolar wall thickness. Kidney sections were evaluated by the following parameters: capillary congestion, hemorrhage, edema, inflammatory cell infiltration, tubular dilation, tubular degeneration, and necrosis. Morphological damage of spleen sections was mainly evaluated based on features of cell apoptosis, including cell shrinkage and nuclear condensation. The percentages of these parameters were counted on a scale of 0–10: 0, not present (normal); 1–4, 10–40% (mild); 5–6, 50–60% (moderate); 7–8, 70–80% (severe); 9–10, 90–100% (very severe). Histological scoring was conducted by an experienced pathologist blinded to genotype and treatment.

**Statistics and reproducibility**

For most experiments, comparisons were made using a 2-tailed, unpaired, Student's t-test. Values shown in each figure represent mean ± SD. A $p\text{-value}<0.05$ was considered statistically significant. Unless otherwise specified, results are representative of at least three independent experiments and means are of at least three individual replicates. For *in vivo* survival experiments, Kaplan-Meier survival curves were performed. All statistical analyses and graphics were made using GraphPad Prism (GraphPad, San Diego, CA).

#### Supplementary Text

Many proteins undergo post-translational modifications (PTMs) to modulate host immune responses. For instance, E3 ubiquitin ligase TRIM31 attenuates inflammasome activation by promoting ubiquitination and proteasomal degradation of NLRP3, while MAVS and E3 ligase TRAF3 promote inflammasome activation by targeting ASC for K63-linked ubiquitination(*5*-*9*). Gasdermin B, a member of the GSDM family, is also modified by ubiquitinoylation and regulates pyroptosis in NK cells(*10*). A recent study showed that poly-ubiquitinoylation of GSDMD at K203/K204 promotes GSDMD-induced pyroptotic cell death(*11*). S-palmitoylation controls numerous biological processes such membrane localization and protein stability(*12*-*14*). MYD88 can be S-palmitoylated, regulating inflammation(*15*). Other proteins such as NOD-1/2 can also be S-palmitoylated, which plays a role in bacterial sensing and membrane localization.(*16*) Palmitoylation is reversible and can regulate other potential PTMs such as phosphorylation(*17*); for example, the palmitoylation/de-palmitoylation cycle promotes STAT3 phosphorylation and activity(*13*). GSDMD is crucial for pyroptosis and plays a critical role in infection and inflammation, and its inhibition protects mice from lethal sepsis(*18*-*20*). Here we identified palmitoylation of GSDSD on Cys191/192 as a regulatory mechanism controlling GSDMD membrane localization and activation, providing a novel target for modulating immune activity in infectious and inflammatory diseases.

GSDMD interacted with FASN, which was essential for its palmitoylation. FASN controls numerous biological functions, including NLRP3 and IL-1β expression, in macrophages(*21*). Previous studies have shown that FASN deficiency in mice attenuates inflammatory responses. Here we reveal that silencing FASN significantly reduced GSDMD palmitoylation and suppressed macrophage pyroptosis. The intracellular dynamics of palmitoylation are tightly regulated by the ZDHHC enzyme family. In human macrophage-like THP-1 cells, GSDMD required at least two palmitoyl acyltransferases, ZDHH5 and ZDHHC9, for its palmitoylation; silencing them by siRNA reduced GSDMD palmitoylation and pyroptotic cell death. Several ZDHHC5/9-mediated palmitoylation events have been reported. ZDHHC5 palmitoylates NOD1/2(*16*), contributing to bacterial sensing. Other ZDHHC5 targets include Flotillins, gp130, cd36, STAT3, S1PR1, and CTNND2(*13, 22*-*26*). ZDHHC9 regulates palmitoylation of a different set of proteins, including HRAS, NRAS, and certain GPCRs(*27*-*29*). It is intriguing that GSDMD required more than one enzyme for its full palmitoylation in human macrophages. Knocking down ZDHHC5 and 9 individually led to 70% and 60% reductions in GSDMD-NT palmitoylation but knocking them both down led to almost complete inhibition (**Fig. 4**). These findings highlight the importance of

GSDMD palmitoylation by ZDHHC5/9 as a potential and novel therapeutic option for treating sepsis and other inflammatory diseases.

GSDMD was palmitoylated at Cys191/192, a residue crucial for GSDMD oligomerization and pore formation(*4*). Disruption of Cys191/192 by site-directed mutagenesis suppresses pyroptotic cell death, and many GSDMD inhibitors exert their function by directly targeting Cys191/192. For example, NSA binds to GSDMD at Cys191/192, abolishing cell death in both macrophages and HEK-293T cells expressing GSDMD-NT and protecting mice from lethal sepsis(*18*). NSA inhibits oligomerization of GSDMD dimers but not GSDMD cleavage nor its initial dimerization. Disulfiram, an FDA-approved drug for alcohol addiction, also suppresses LPS-induced septic death in mice by covalently modifying GSDMD Cys191/Cys192 to block pore formation, pyroptosis, and IL-1β release(*20*). Similarly, treatment with dimethyl fumarate (DMF), a fumarate derivative and tricarboxylic cycle intermediate, abolishes cell death in both human and mouse macrophages upon LPS/Nig stimulation. DMF reacts with GSDMD Cys191/Cys192 to form S-(2-succinyl)-cysteine, a PTM known as succination, which prevents GSDMD-caspase interactions to limit GSDMD processing, oligomerization, and capacity to induce cell death(*19*). Administration of DMF protects mice from LPS-induced sepsis and reduces the severity of experimental allergic encephalitis by targeting GSDMD(*19*), reinforcing the importance of Cys191/192 in regulating GSDMD activity. GSDMD Cys191/Cys192 has also been reported to act as a pyroptosis-inducing redox sensor(*30, 31*). The Ragulator-Rag-mTORC1 pathway, by mediating ROS production, promotes pyroptotic cell death. Diverse innate immune signaling pathways or environmental perturbations that induce ROS appear to bypass the requirement for Ragulator-Rag. ROS facilitate NT-GSDMD oligomerization and pore formation, probably through oxidative modification of cysteine residues, providing another checkpoint for pyroptosis and suggesting that there are key biochemical steps for efficient pore formation even after GSDMD cleavage(*30, 31*). Although GSDMD can be directly oxidized at multiple cysteine residues, only Cys191/192 is required for ROS-mediated potentiation of GSDMD pore formation and pyroptosis. The precise mechanism of how cysteine oxidation within GSDMD contributes to pore formation has yet to be defined. Since the same Cys191/192 can be robustly palmitoylated, the mechanisms already thought to be mediated by Cys191/192 and many Cys191/192-mediated drug effects might potentially be regulated by palmitoylation.

Both full-length GSDMD and GSDMD-NT could be palmitoylated. Intriguingly, palmitoylated full-length GSDMD remained cytosolic, suggesting that the palmitoyl moiety might be sequestered in a hydrophobic cleft within the full-length protein, and therefore might not be exposed to promote membrane binding. In macrophages, palmitoylation of full-length GSDMD was also triggered by LPS stimulation. Thus, priming with LPS can prepare GSDMD for membrane localization even before its cleavage by caspase 1/11, ensuring efficient GSDMD activation and pore-forming activity on the membrane after caspase 1/11 activation. Of note, a recent study revealed a conserved pore-forming domain in bacterial gasdermins that is stabilized in the inactive state with a buried palmitoyl modification(*32*). Presumably, S-palmitoylation may also contribute to stability of the inactive mammalian full-length GSDMD. In another study, GSDME was found to be palmitoylated on its C-terminal (GSDME-CT) during chemotherapy-induced cancer cell pyroptosis. This palmitoylation event blocks the interaction between GSDME-NT and GSDME-CT, facilitates their dissociation, and thus promotes GSDME-NT pore forming activity(*33*). Palmitoylation at GSDMD-NT may deliver the same effect; however, palmitoylation itself is not able to overcome the autoinhibition by the GSDMD-CT in the absence of caspase cleavage.

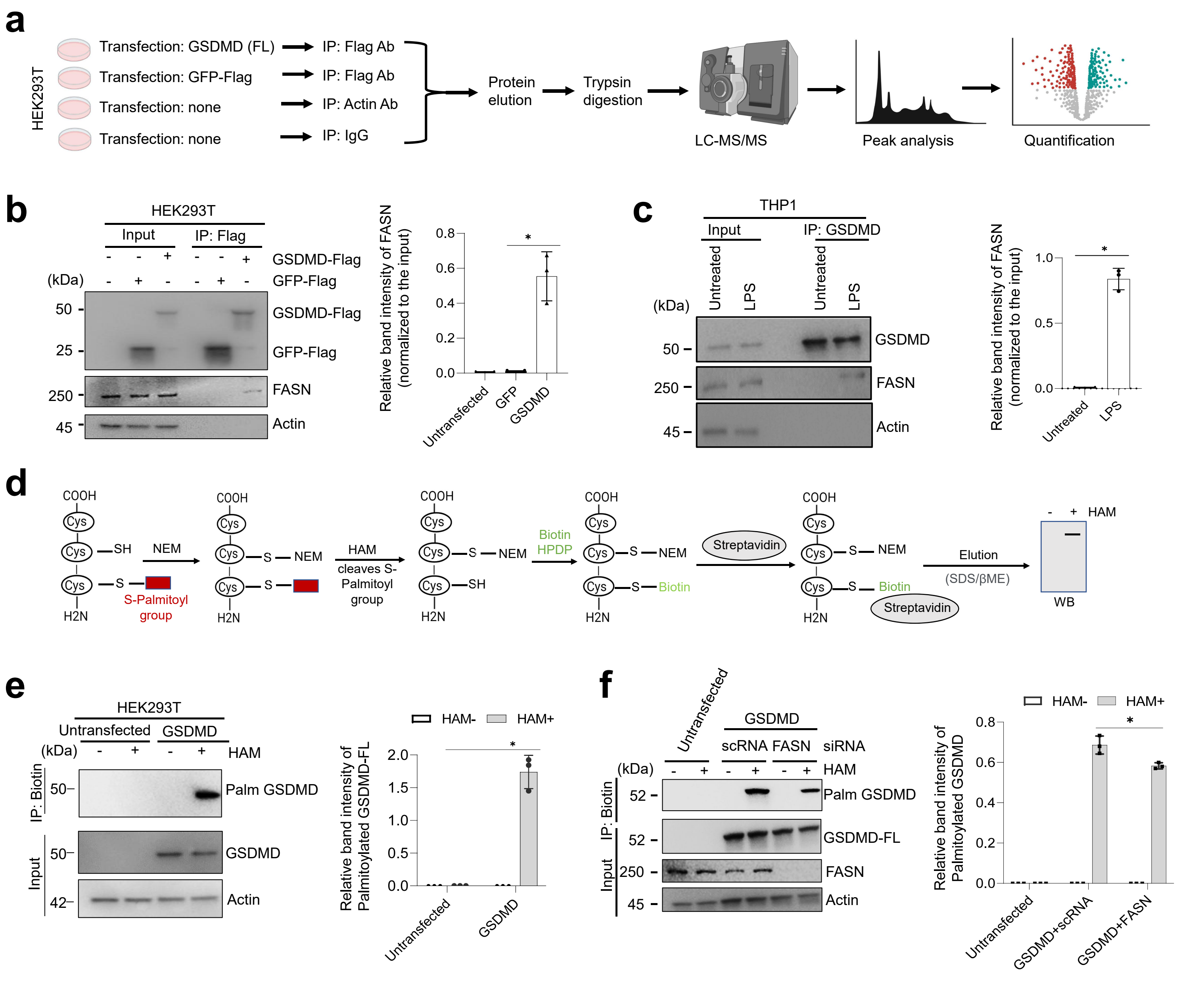
a
HEK293T
Transfection: GSDMD (FL)
Transfection: GFP-Flag
Transfection: none
Transfection: none
IP: Flag Ab
IP: Flag Ab
IP: Actin Ab
IP: IgG
Protein elution
Trypsin digestion
LC-MS/MS
Peak analysis
Quantification
b
HEK293T
Input
IP: Flag
GSDMD-Flag
GFP-Flag
(kDa)
50
25
250
45
FASN
Actin
Relative band intensity of FASN (normalized to the input)
Untransfected
GFP
GSDMD
c
THP1
Input
IP: GSDMD
Untreated
LPS
GSDMD
FASN
Actin
d
COOH
Cys
SH
NEM
S-Palmitoyl group
H2N
HAM cleaves S-Palmitoyl group
Biotin HPDP
Biotin
Streptavidin
Elution
(SDS/βME)
HAM
WB
e
HEK293T
Untransfected
GSDMD
HAM
IP: Biotin
Input
Palm GSDMD
GSDMD
Actin
HAM-
HAM+
Relative band intensity of Palmitoylated GSDMD-FL
f
Untransfected
GSDMD
scRNA FASN siRNA
Palm GSDMD
GSDMD-FL
FASN
Actin
Relative band intensity of Palmitoylated GSDMD
GSDMD+scRNA
GSDMD+FASN

**Fig. S1. GSDMD interacts with FASN**

(a) Schematic of the approach and strategy used to identify GSDMD-interacting proteins by mass spectrometry.

(b) HEK293T cells transfected with or without hGSDMD-FL-FLAG were co-immunoprecipitated with anti-FLAG antibodies, and immunoblot analyses were performed with anti-FLAG HRP antibodies. Right, quantification of relative FASN protein bound with GSDMD-FL.

(c) PMA-differentiated THP-1 cells were treated with or without LPS (1 μg/ml) for 3 h. Cells were lysed, and immunoprecipitation was performed with anti-rabbit GSDMD-FL antibody followed by western blot analysis. Right, image represents the quantification of endogenous FASN bound with GSDMD protein.

(d) Schematic of the steps in the acyl-biotin exchange (ABE) palmitoylation assay.

(e) HEK293T cells were transfected with or without GSDMD-FL. After 24 h, cell lysates were lysed and blocked with NEM followed by hydroxylamine treatment. Proteins were then reacted with biotin-HPDP and the reacted proteins pulled down with streptavidin beads. The eluted proteins with or without hydroxylamine were probed with anti-FLAG HRP antibody. Right, quantification of palmitoylated GSDMD proteins normalized to input.

(f) FLAG-hGSDMD-FL was co-expressed with control or *FASN* siRNA and subjected to the ABE palmitoylation assay and immunoblot analyses. Right, quantification of palmitoylated GSDMD proteins normalized to input.

Data are mean ± SEM (b-c, e-f). Western blots shown are representative of three independent experiments. *$P<0.05$, **$P<0.01$, ***$P<0.001$, unpaired Student's *t*-test.

Fig.S2

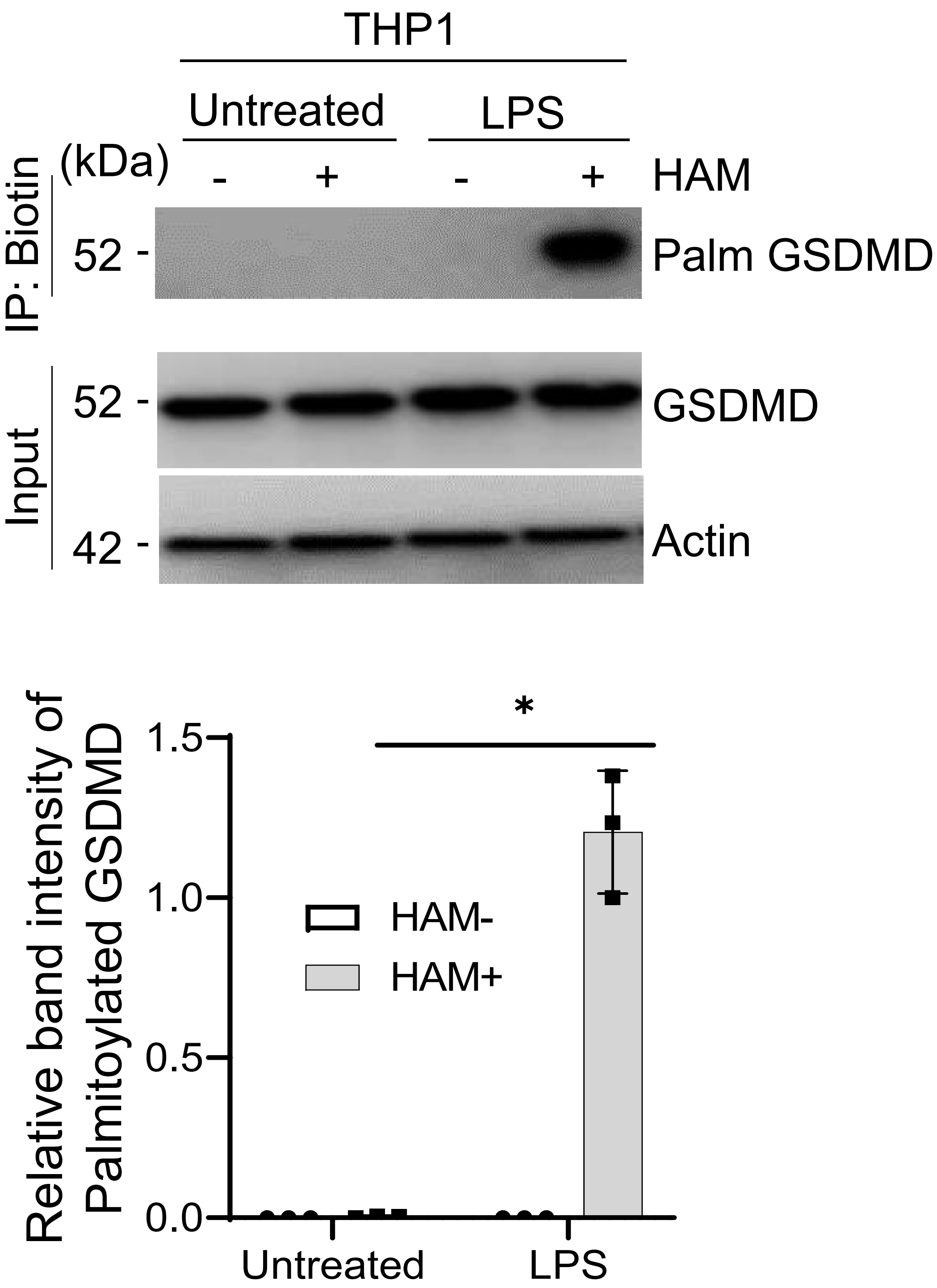
THP1
Untreated
LPS
(kDa)
- + - + HAM
IP: Biotin
52 -
Palm GSDMD
Input
52 -
GSDMD
42 -
Actin
*
1.5
1.0
0.5
0.0
HAM-
HAM+
Relative band intensity of Palmitoylated GSDMD
Untreated
LPS

**Fig. S2. LPS triggers GSDMD palmitoylation in macrophages.**

THP-1 cells treated with or without LPS (3 h) were subjected to the ABE assay followed by the presence or absence of hydroxylamine. Immunoblot analyses were performed with anti-GSDMD antibodies. Western blot images are representative of three independent experiments. ***$p<0.001$, unpaired two-tailed Student's *t*-test.

Fig.S3

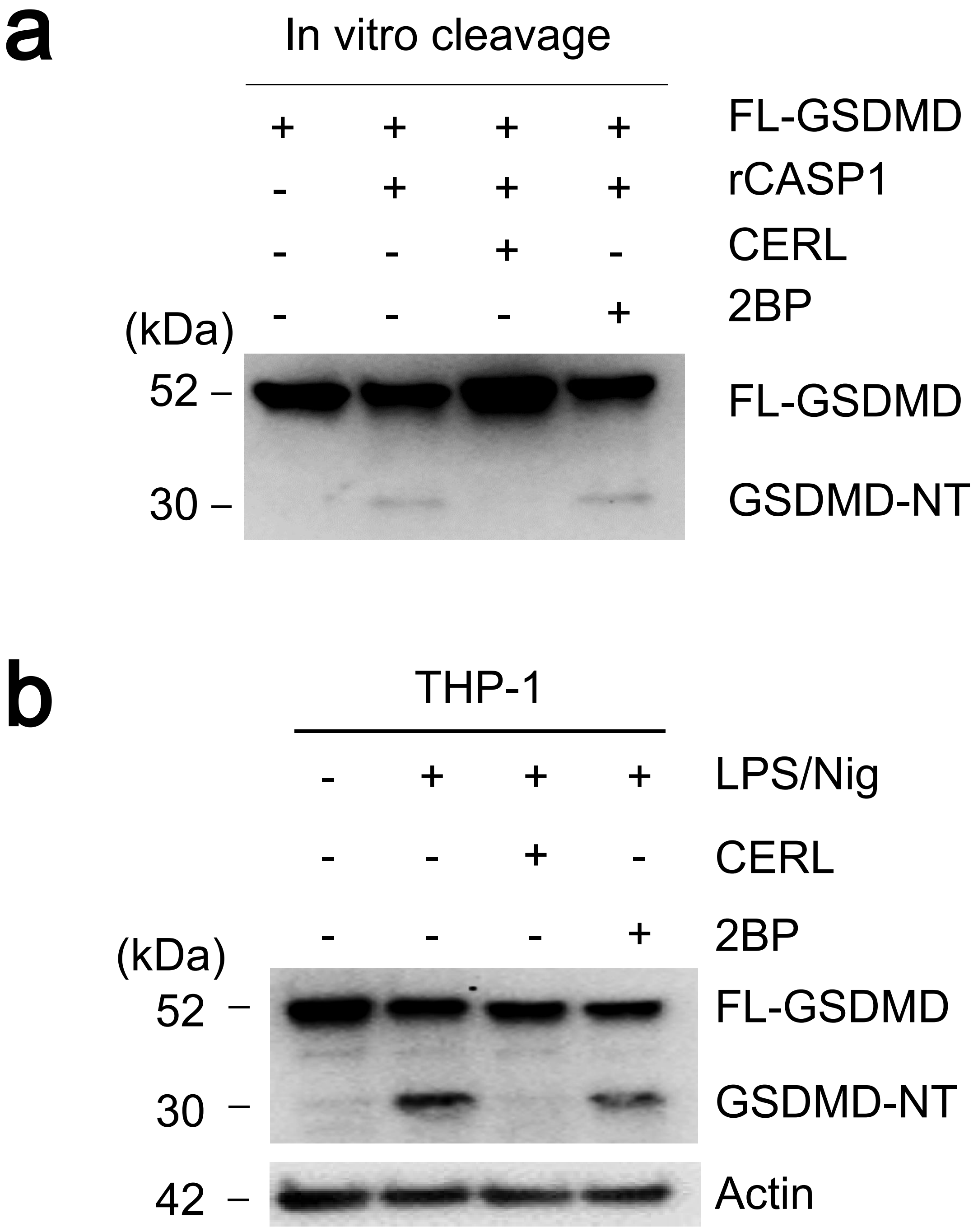
a
In vitro cleavage
FL-GSDMD
rCASP1
CERL
2BP
(kDa)
52
30
FL-GSDMD
GSDMD-NT
b
THP-1
LPS/Nig
CERL
2BP
(kDa)
52
30
42
FL-GSDMD
GSDMD-NT
Actin

**Fig. S3. Palmitoylation is not required for GSDMD cleavage.**

(a) *In vitro* cleavage assay of recombinant GSDMD-FL treated with or without active caspase-1 in the presence or absence of palmitoylation inhibitors. Immunoblots were performed with anti-GSDMD antibodies.

(b) THP-1 cells primed with LPS for 3 h followed by treatment with palmitoylation inhibitors (cerulenin, 50 μM; 2BP, 10 μM) for 30 mins before stimulation with nigericin for 35 min. Immunoblots were performed with anti-GSDMD-FL antibodies. Western blot images are representative of three independent repeats.

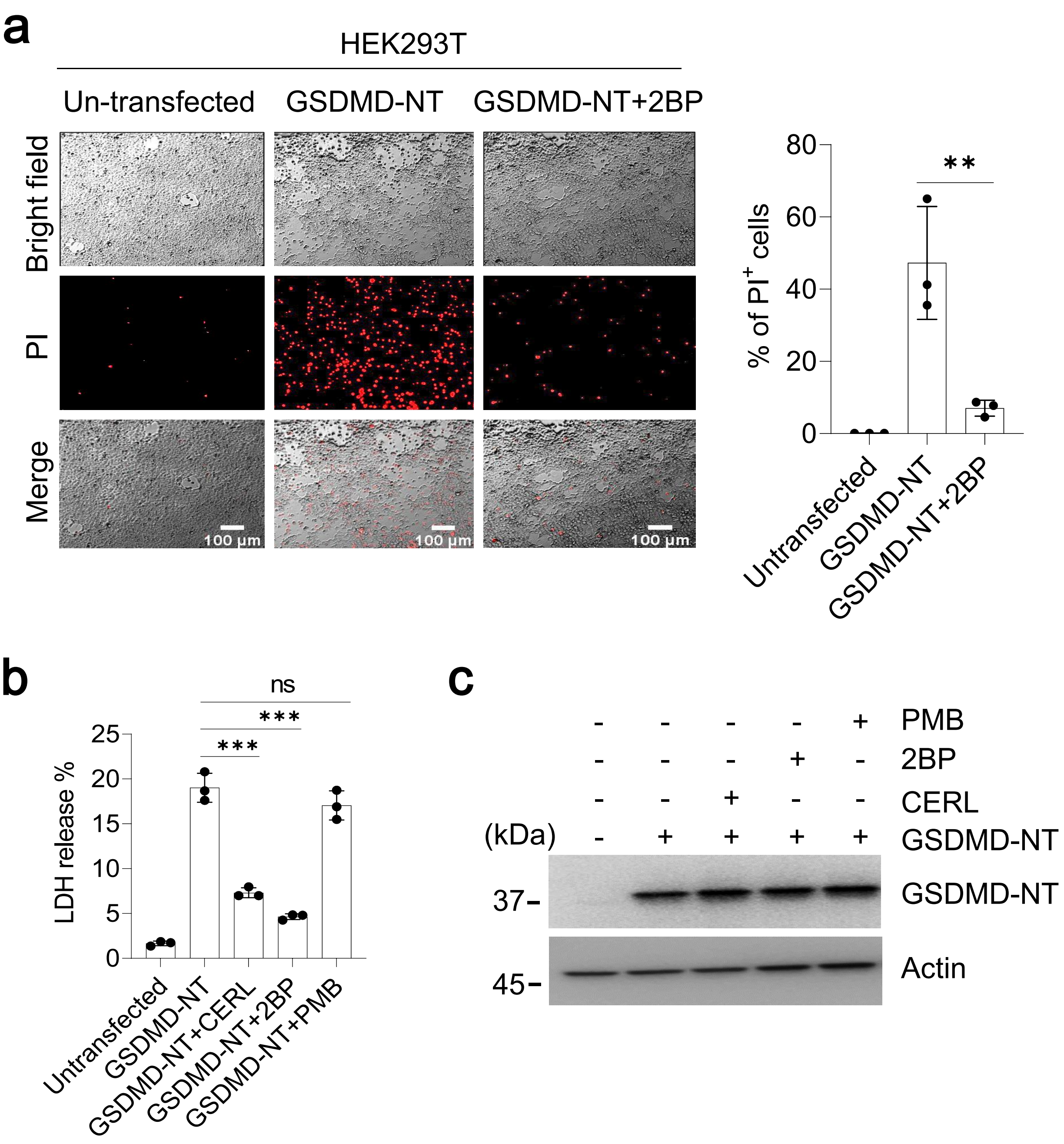
a
HEK293T
Un-transfected
GSDMD-NT
GSDMD-NT+2BP
Bright field
PI
Merge
100 μm
100 μm
100 μm
% of PI+ cells
80
60
40
20
0
**
Untransfected
GSDMD-NT
GSDMD-NT+2BP
b
LDH release %
25
20
15
10
5
0
ns
***
***
Untransfected
GSDMD-NT
GSDMD-NT+CERL
GSDMD-NT+2BP
GSDMD-NT+PMB
c
PMB
2BP
CERL
(kDa)
GSDMD-NT
37-
GSDMD-NT
45-
Actin

**Fig. S4. Palmitoylation inhibitors reduce GSDMD-NT pyroptotic activity in 293T cells**.

(a) 293T cells transfected with or without GSDMD-NT were treated with or without 2BP (50 µM). Cells were stained with PI and fluorescence images captured. Right, image represents the quantification of relative PI uptake in 293T cells.

(b) LDH release from cell culture supernatants of 293T cells treated with or without palmitoylation inhibitors.

(c) Western blot images of 293T cells showing the expression of GSDMD-NT. All data are representative of three independent experiments.

Data are mean ± SEM. $^{**}P<0.01$, $^{***}P<0.001$, unpaired two-tailed Student's *t*-test.

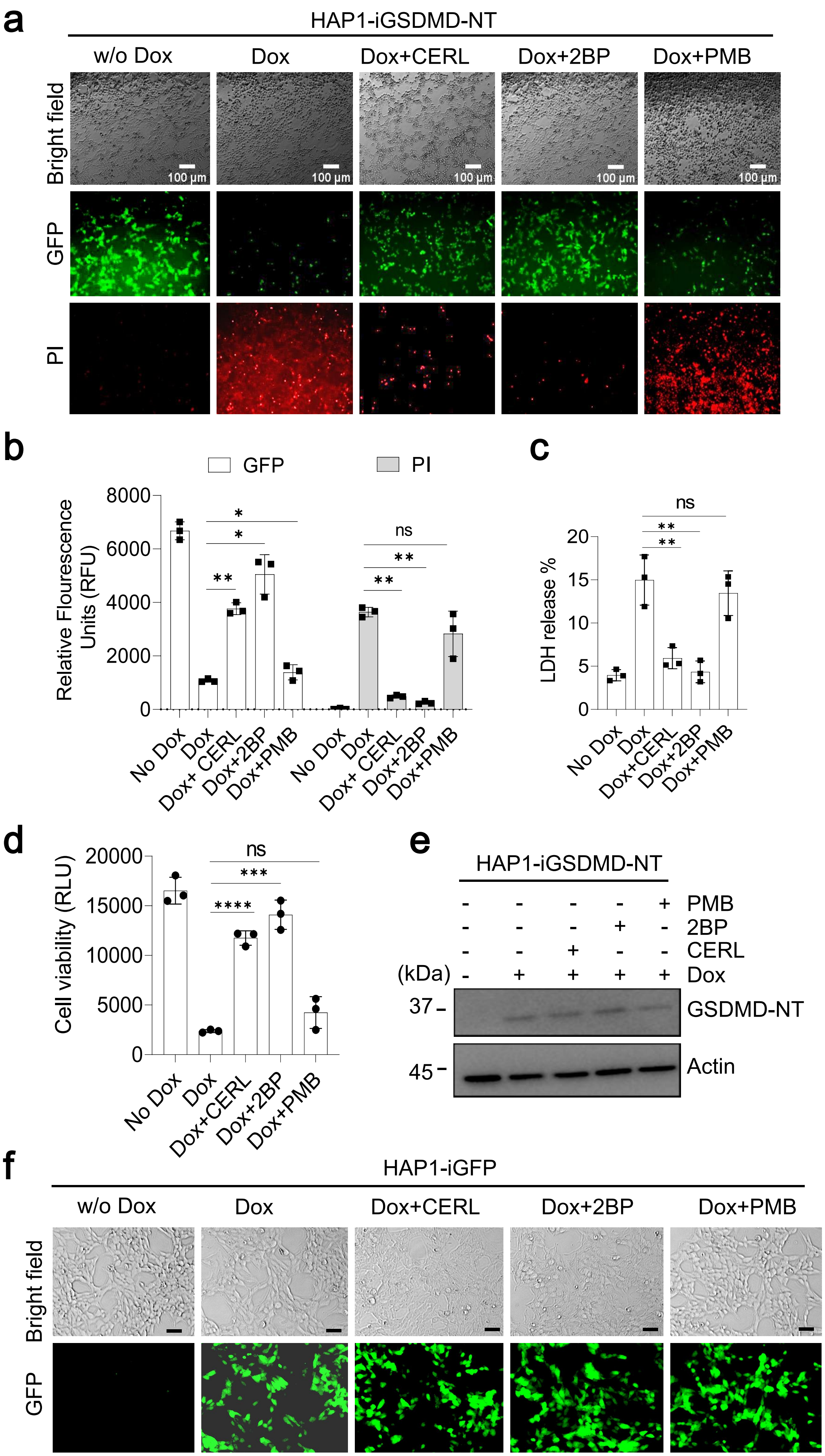
a
HAP1-iGSDMD-NT
w/o Dox
Dox
Dox+CERL
Dox+2BP
Dox+PMB
Bright field
GFP
PI
100 µm
b
GFP
PI
Relative Flourescence Units (RFU)
8000
6000
4000
2000
0
ns
No Dox
Dox
Dox+ CERL
Dox+2BP
Dox+PMB
c
ns
LDH release %
20
15
10
5
0
No Dox
Dox
Dox+CERL
Dox+2BP
Dox+PMB
d
ns
Cell viability (RLU)
20000
15000
10000
5000
0
No Dox
Dox
Dox+CERL
Dox+2BP
Dox+PMB
e
HAP1-iGSDMD-NT
PMB
2BP
CERL
(kDa)
Dox
37
45
GSDMD-NT
Actin
f
HAP1-iGFP
w/o Dox
Dox
Dox+CERL
Dox+2BP
Dox+PMB
Bright field
GFP

**Fig. S5. Palmitoylation is required for pyroptotic cell death directly induced by GSDMD-NT**

(a) To bypass all the upstream molecules inducing pyroptosis, a cleaved N-terminal fragment product and constituently active GFP were cloned into a Tet3G transactivator under a Dox-inducible promoter. HAP1 cells were transduced with GSDMD-NT vector and cells were stably transfected, selected with blasticidin, and single cells were isolated and screened for GSDMD expression. To determine whether GSDMD-NT can induce pore formation after Dox treatment, Dox-induced GSDMD-NT expressing HAP1 cells (**HAP1-iGSDMD-NT**) were primed with palmitoylation inhibitors (cerulenin, 50 μM; 2BP, 50 μM) and a de-palmitoylation inhibitor (palmostatin B 50 μM) for 30 min before treatment with doxycycline (100 ng/ml) for 3 h. Cell dye propidium iodide was added and PI fluorescence captured by fluorescence microscopy. Scale bars, 100 μM.

(b) The relative mean fluorescence intensity of PI and GFP were quantified using ImageJ.

(c) Control and treated cell culture media supernatants were subjected to the LDH assay and LDH percentage release was calculated for each group.

(d) The cells were subjected to a cell viability assay using the RealTime-Glo MT Cell Viability Assay kit (Promega). Relative luminescence units (RLU) were captured with a luminometer.

(e) Western blot images shown are representative blots of doxycycline-inducible HAP1 cells expressing GSDMD-NT (HAP1-iGSDMD-NT). All data are representative of three independent experiments.

Data are mean ± SEM. *P<0.05, **P<0.01, ***P<0.001, unpaired Student's *t*-test.

(f) Inhibition of palmitoylation does not alter GSDMD-NT expression. Fluorescence images of Dox-induced GFP cells treated with or without palmitoylation inhibitors (cerulenin, 50 μM; 2BP, 50 μM) and de-palmitoylation inhibitor palmostatin B (50 μM) for 30 min before treatment with doxycycline (100 ng/ml) for 6 h. Brightfield and GFP signals were captured, and images were taken by fluorescence microscopy. All data are representative of three independent experiments. Scale bars, 100 μM.

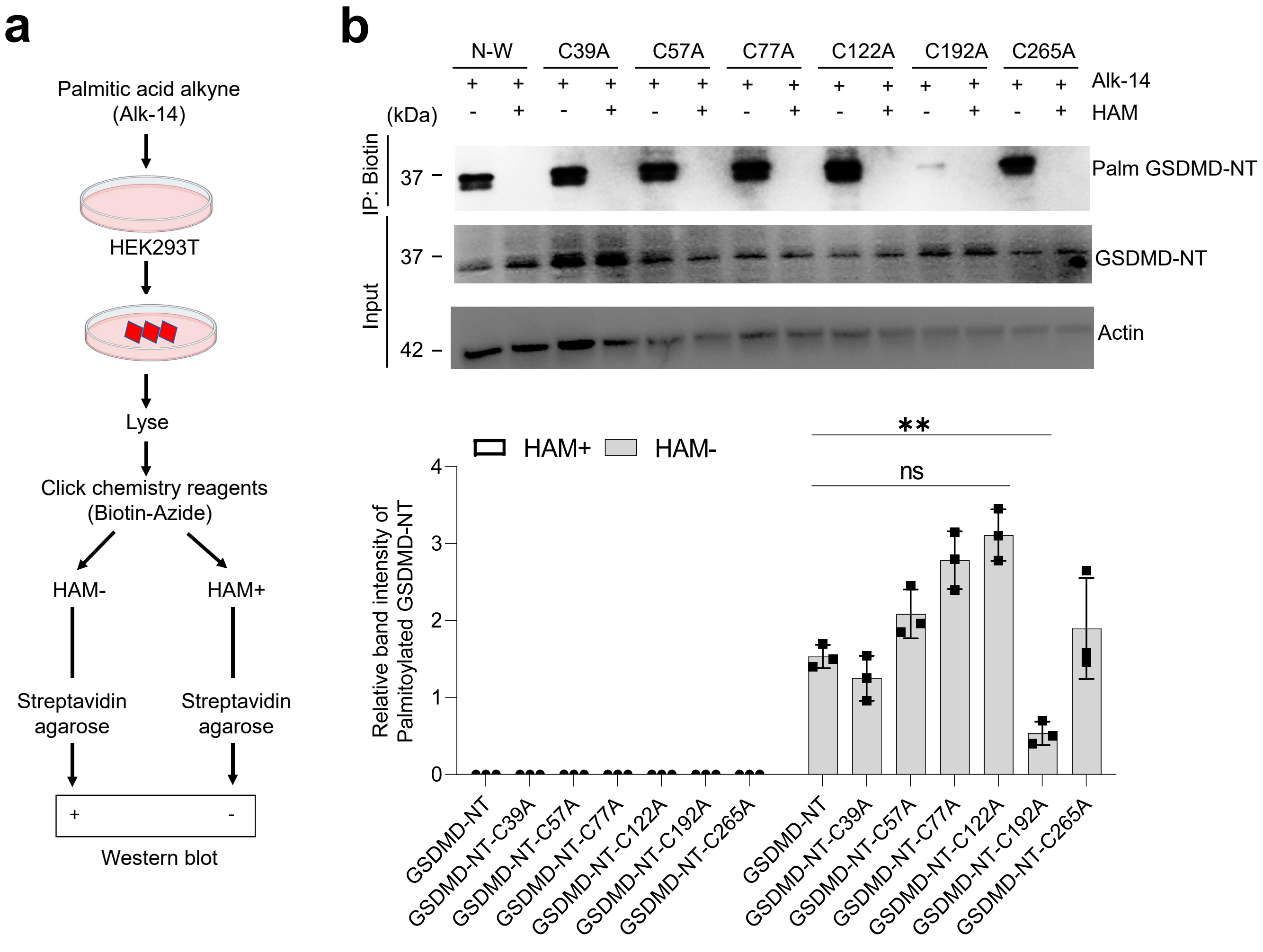
a
Palmitic acid alkyne
(Alk-14)
HEK293T
Lyse
Click chemistry reagents
(Biotin-Azide)
HAM-
HAM+
Streptavidin agarose
Streptavidin agarose
+
-
Western blot
b
N-W
C39A
C57A
C77A
C122A
C192A
C265A
Alk-14
HAM
(kDa)
IP: Biotin
Input
37
37
42
Palm GSDMD-NT
GSDMD-NT
Actin
HAM+
HAM-
**
ns
Relative band intensity of Palmitoylated GSDMD-NT
GSDMD-NT
GSDMD-NT-C39A
GSDMD-NT-C57A
GSDMD-NT-C77A
GSDMD-NT-C122A
GSDMD-NT-C192A
GSDMD-NT-C265A

**Fig. S6. Cys192 is the major palmitoylation site in GSDMD-NT.**

(a) A schematic illustrating the sequential steps involved in palmitic acid labeling to detect the palmitoylation in 293T cells.

(b) 293T cells expressing wild type or mutant GSDMD-NT (Cys39A, Cys57A, Cys77A, Cys122A, Cys192A, Cys265A) were labeled with alkyne-tagged palmitic acid. After 24 h, both cells and supernatant were precipitated with methanol:chloroform, and samples were subjected to click chemistry reagents (biotin azide) in the presence or absence of hydroxylamine. Immunoblot analyses were performed with anti-FLAG antibody. The relative amount of palmitoylated GSDMD-NT was quantified relative to the input. Data are representative of three independent replicates. Data are ± SEM. **$P<0.01$, unpaired two-tailed Student's *t*-test.

.

Fig.S7

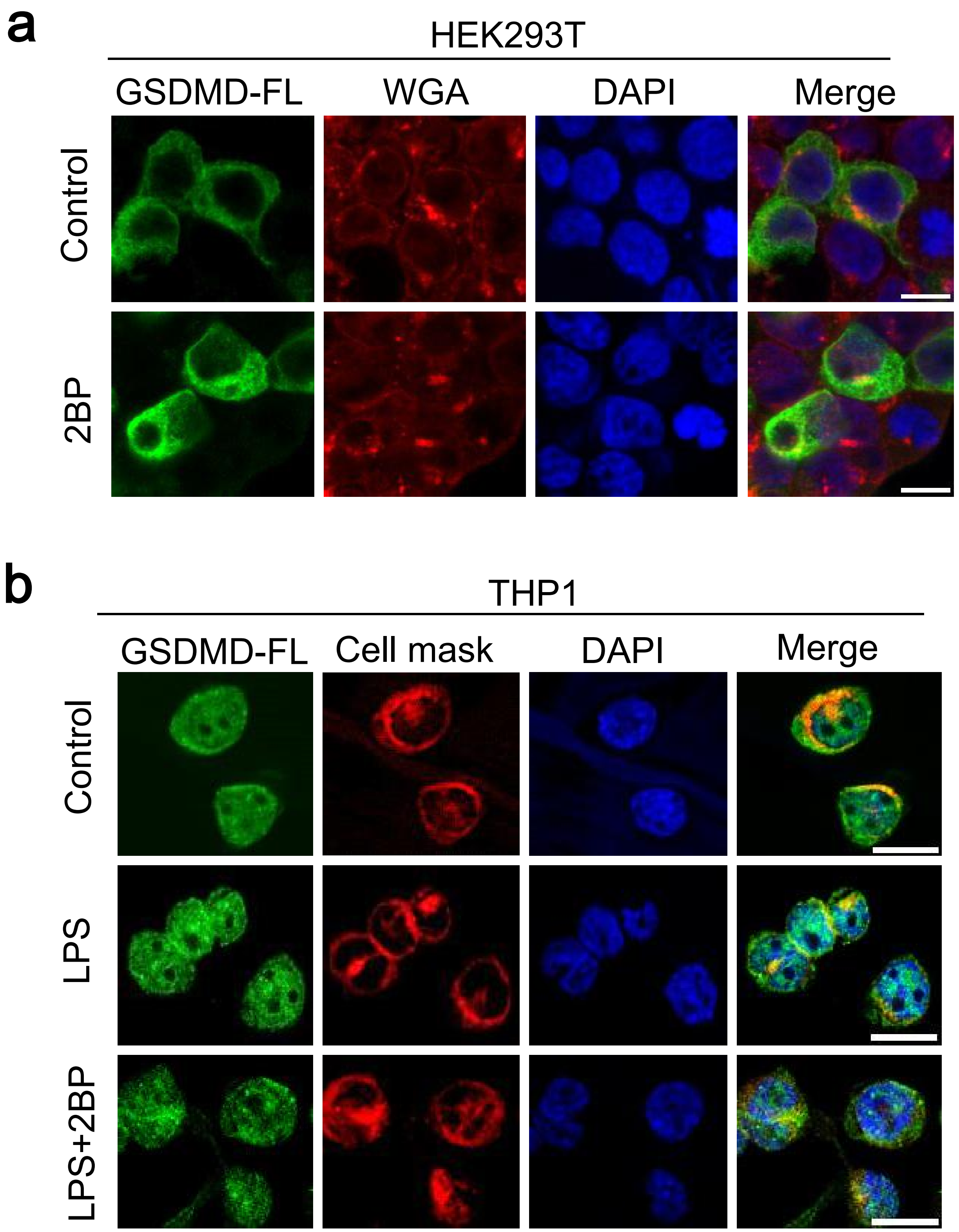
a
HEK293T
GSDMD-FL
WGA
DAPI
Merge
Control
2BP
b
THP1
GSDMD-FL
Cell mask
DAPI
Merge
Control
LPS
LPS+2BP

**Fig. S7. Palmitoylation of GSDMD-FL does not affect its translocation.**

(a) Subcellular localization analysis of 293T cells overexpressing GSDMD-FL treated with or without 2BP by immunofluorescence analysis. Images were taken by confocal microscopy. Scale bars, 20 μm. Images are representative of three independent replicates.

(b) Representative images of subcellular localization in THP-1 cells expressing endogenous GSDMD-FL primed with 2BP and treated with LPS for 3 h. Images were analyzed by confocal microscopy. Scale bars, 20 μm.

Fig.S8

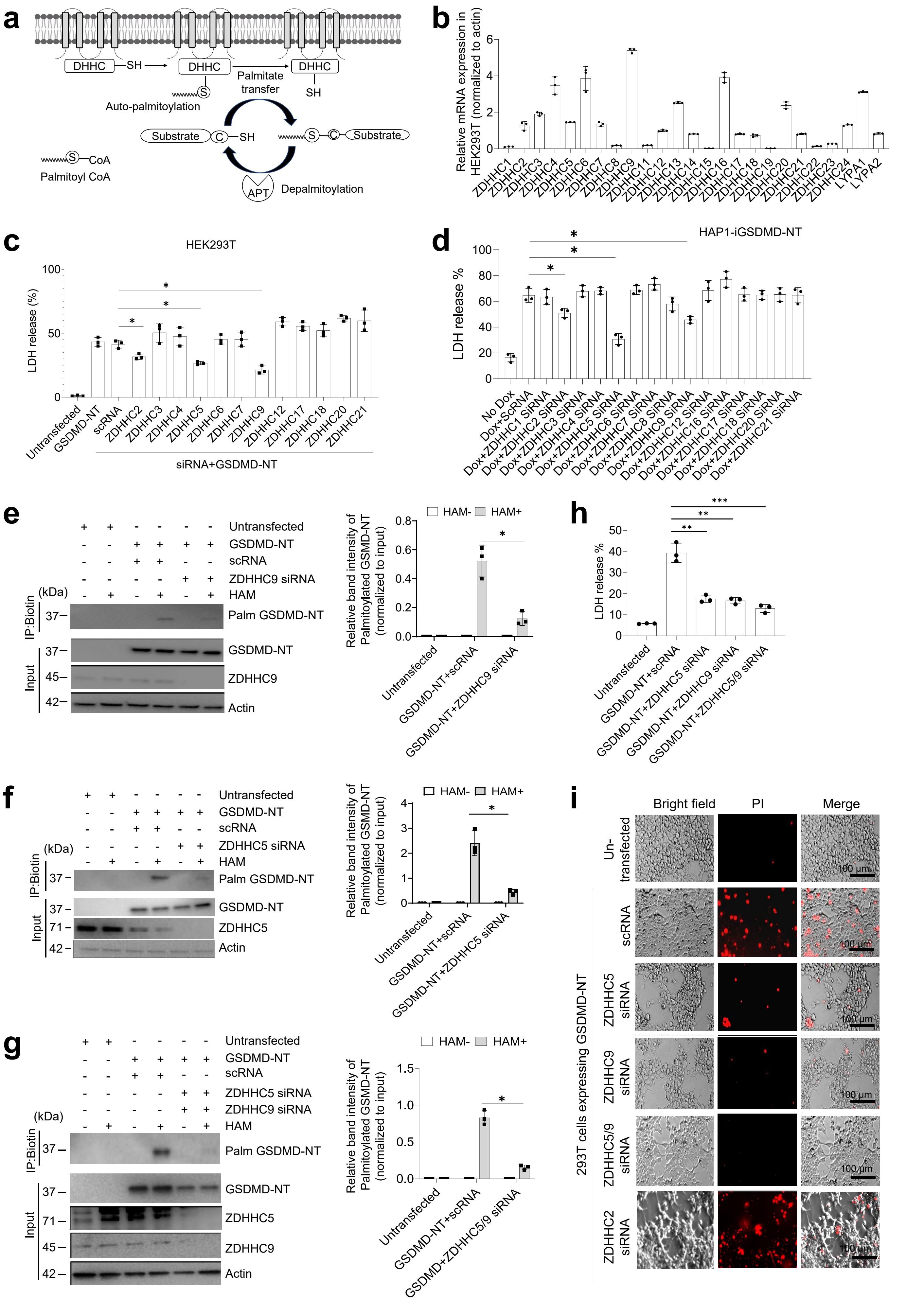
a
DHHC
SH
DHHC
DHHC
Palmitate transfer
SH
S
Auto-palmitoylation
Substrate
C
SH
S
C
Substrate
S
CoA
Palmitoyl CoA
APT
Depalmitoylation
b
Relative mRNA expression in HEK293T (normalized to actin)
ZDHHC1
ZDHHC2
ZDHHC3
ZDHHC4
ZDHHC5
ZDHHC6
ZDHHC7
ZDHHC8
ZDHHC9
ZDHHC11
ZDHHC12
ZDHHC13
ZDHHC14
ZDHHC15
ZDHHC16
ZDHHC17
ZDHHC18
ZDHHC19
ZDHHC20
ZDHHC21
ZDHHC22
ZDHHC23
ZDHHC24
LYPA1
LYPA2
c
HEK293T
LDH release (%)
Untransfected
GSDMD-NT
scRNA
ZDHHC2
ZDHHC3
ZDHHC4
ZDHHC5
ZDHHC6
ZDHHC7
ZDHHC9
ZDHHC12
ZDHHC17
ZDHHC18
ZDHHC20
ZDHHC21
siRNA+GSDMD-NT
d
HAP1-iGSDMD-NT
LDH release %
No Dox
Dox+ScRNA
Dox+ZDHHC1 SiRNA
Dox+ZDHHC2 SiRNA
Dox+ZDHHC3 SiRNA
Dox+ZDHHC4 SiRNA
Dox+ZDHHC5 SiRNA
Dox+ZDHHC6 SiRNA
Dox+ZDHHC7 SiRNA
Dox+ZDHHC8 SiRNA
Dox+ZDHHC9 SiRNA
Dox+ZDHHC12 SiRNA
Dox+ZDHHC16 SiRNA
Dox+ZDHHC17 SiRNA
Dox+ZDHHC18 SiRNA
Dox+ZDHHC20 SiRNA
Dox+ZDHHC21 SiRNA
e
Untransfected
GSDMD-NT
scRNA
ZDHHC9 siRNA
(kDa)
HAM
IP:Biotin
Palm GSDMD-NT
Input
GSDMD-NT
ZDHHC9
Actin
HAM-
HAM+
Relative band intensity of Palmitoylated GSMD-NT (normalized to input)
Untransfected
GSDMD-NT+scRNA
GSDMD-NT+ZDHHC9 siRNA
h
LDH release %
Untransfected
GSDMD-NT+scRNA
GSDMD-NT+ZDHHC5 siRNA
GSDMD-NT+ZDHHC9 siRNA
GSDMD-NT+ZDHHC5/9 siRNA
f
Untransfected
GSDMD-NT
scRNA
ZDHHC5 siRNA
(kDa)
HAM
IP:Biotin
Palm GSDMD-NT
Input
GSDMD-NT
ZDHHC5
Actin
HAM-
HAM+
Relative band intensity of Palmitoylated GSMD-NT (normalized to input)
Untransfected
GSDMD-NT+scRNA
GSDMD-NT+ZDHHC5 siRNA
i
Bright field
PI
Merge
Un-transfected
293T cells expressing GSDMD-NT
scRNA
ZDHHC5 siRNA
ZDHHC9 siRNA
ZDHHC5/9 siRNA
ZDHHC2 siRNA
100 μm
g
Untransfected
GSDMD-NT
scRNA
ZDHHC5 siRNA
ZDHHC9 siRNA
(kDa)
HAM
IP:Biotin
Palm GSDMD-NT
Input
GSDMD-NT
ZDHHC5
ZDHHC9
Actin
HAM-
HAM+
Relative band intensity of Palmitoylated GSMD-NT (normalized to input)
Untransfected
GSDMD-NT+scRNA
GSDMD+ZDHHC5/9 siRNA

**Fig. S8. GSDMD-NT palmitoylation is mediated by ZDHHC5 and ZDHHC9, which are essential for GSDMD-NT-induced pyroptosis**

**a,** Graphical illustration depicts the overall process of DHHC-mediated S-palmitoylation.

**b,** Real time-PCR analysis of ZDHHC expression in 293T cells. The relative mRNA expression was calculated and normalized to actin as an internal control. Data are presented as the mean ± SEM.

**c,** 293T cells were co-transfected with or without GSDMD-NT and *ZDHHC* siRNA for 24 h. The cell culture media were collected and quantified for LDH release. Luminescence was captured with a luminometer.

**d,** HAP1 cells expressing GSDMD-NT (HAP1-iGSDMD-NT) were transfected with control or *ZDHHC* siRNA followed by stimulation with doxycycline for 3 h. LDH release from the supernatant was quantified.

**e-g,** 293T cells transfected with GSDMD-NT and *ZDHHC* siRNA were subjected to the ABE assay and immunoblot analysis performed. Right, image represents the quantification of palmitoylated GSDMD-NT normalized to input.

**h,** LDH release from the culture supernatant of 293T cells was quantified.

**i,** Representative fluorescence images of PI uptake in 293T cells expressing GSDMD-NT and *ZDHHC* siRNA. Scale bars, 100 μM.

Data are presented as the mean ± SEM. Western blot Images shown are representative of three independent experiments. *P<0.05, **P<0.01, ***P<0.001, unpaired Student's *t*-test.

**a**

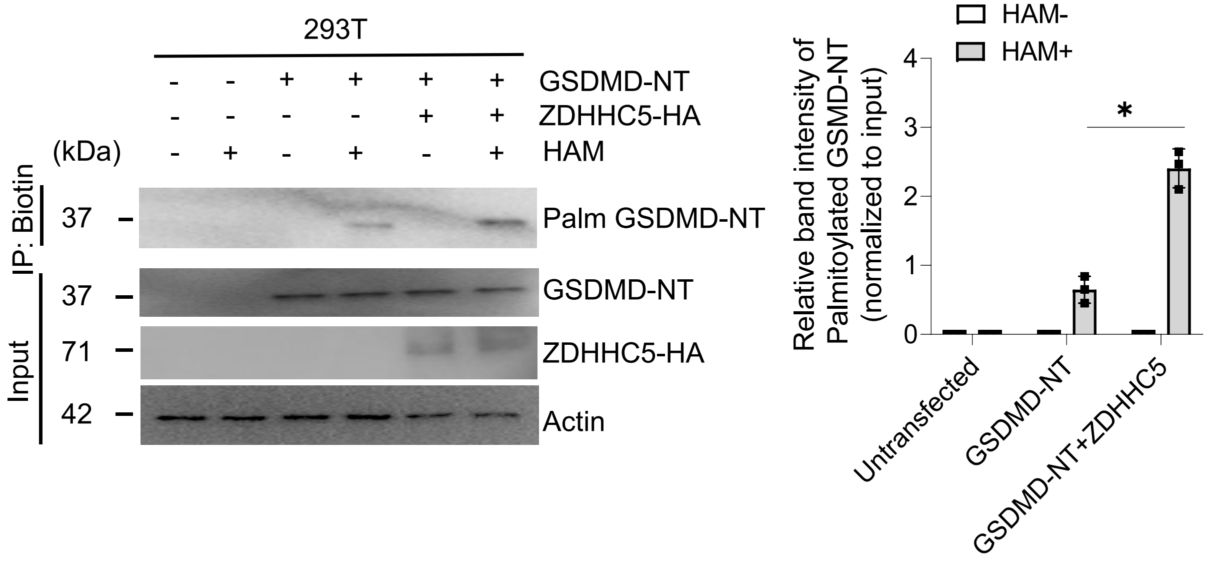
293T
GSDMD-NT
ZDHHC5-HA
HAM
(kDa)
IP: Biotin
Input
37
37
71
42
Palm GSDMD-NT
GSDMD-NT
ZDHHC5-HA
Actin
HAM-
HAM+
Relative band intensity of Palmitoylated GSMD-NT (normalized to input)
*
Untransfected
GSDMD-NT
GSDMD-NT+ZDHHC5

**b**

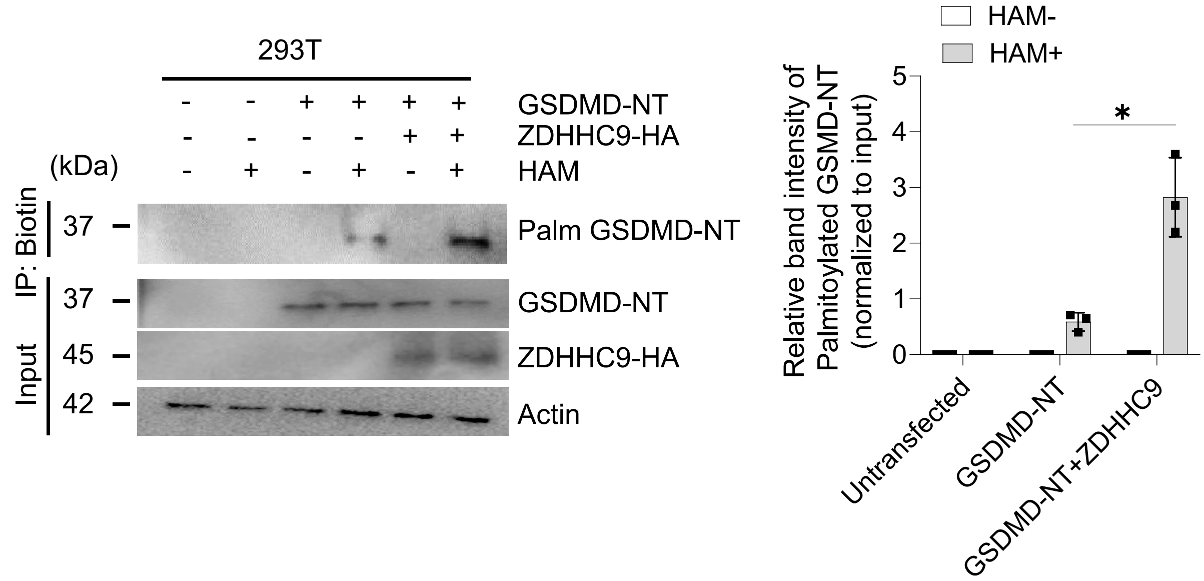
293T
GSDMD-NT
ZDHHC9-HA
HAM
(kDa)
IP: Biotin
Input
37
37
45
42
Palm GSDMD-NT
GSDMD-NT
ZDHHC9-HA
Actin
HAM-
HAM+
Relative band intensity of Palmitoylated GSMD-NT (normalized to input)
*
Untransfected
GSDMD-NT
GSDMD-NT+ZDHHC9

**Fig. S9. Overexpression of ZDHHC5 or ZDHHC9 leads to increased palmitoylation of GSDMD-NT.**

HEK293T cells were co-transfected with GSDMD-NT and HA-tagged ZDHHC5 (a) or ZDHHC9 (b) plasmids and ABE analyses performed. Immunoblot analysis was performed to detect GSDMD-NT palmitoylation. Right, images shown are quantification of palmitoylated GSDMD-NT in 293T cells. Data are representative of three independent replicates. Data are mean ± SEM. **$P<0.01$, unpaired two-tailed Student's *t*-test.

Fig.S10

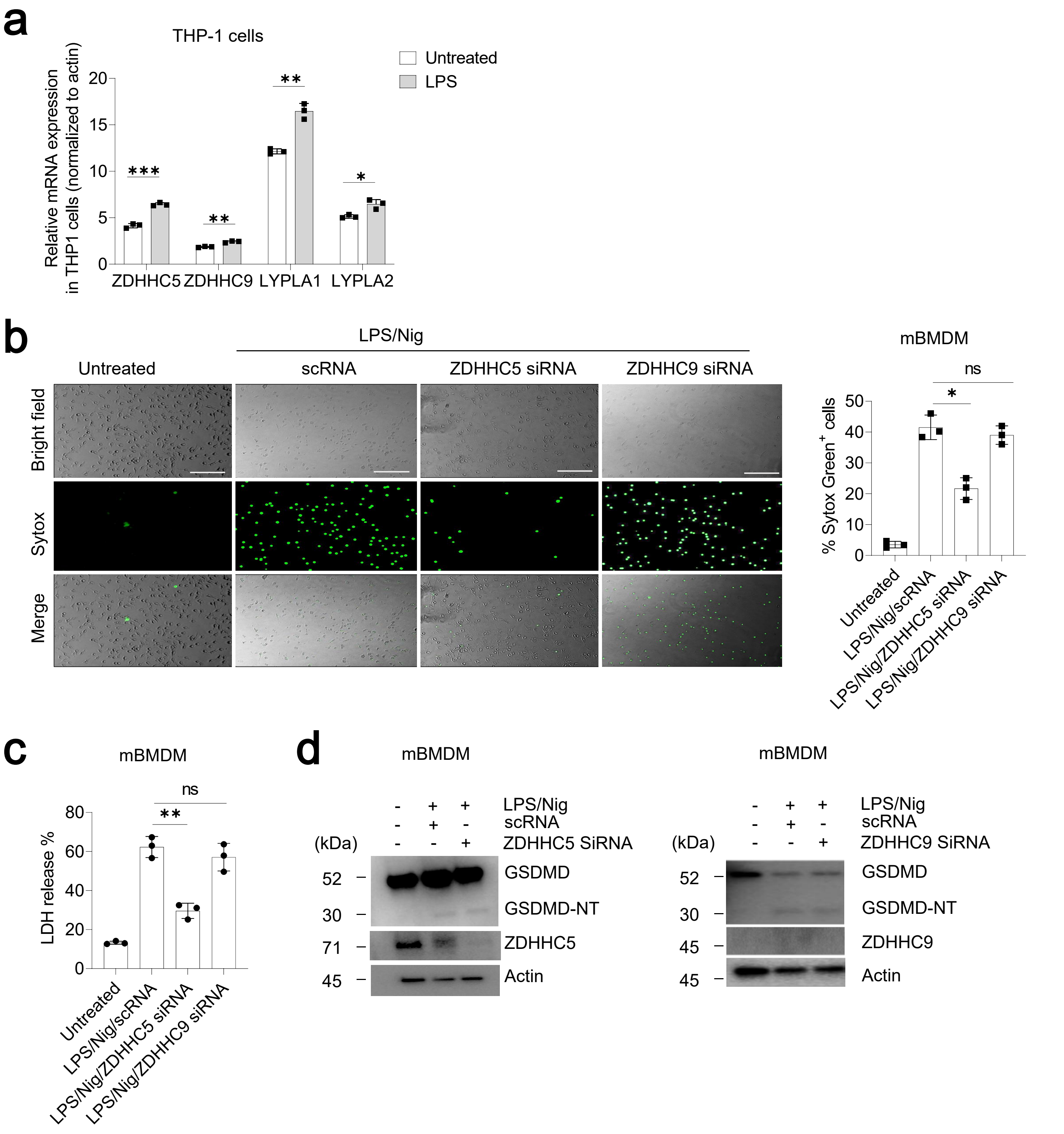
a
THP-1 cells
Untreated
LPS
Relative mRNA expression in THP1 cells (normalized to actin)
20
15
10
5
0
***
**
**
*
ZDHHC5
ZDHHC9
LYPLA1
LYPLA2
b
LPS/Nig
Untreated
scRNA
ZDHHC5 siRNA
ZDHHC9 siRNA
Bright field
Sytox
Merge
mBMDM
ns
*
% Sytox Green+ cells
50
40
30
20
10
0
Untreated
LPS/Nig/scRNA
LPS/Nig/ZDHHC5 siRNA
LPS/Nig/ZDHHC9 siRNA
c
mBMDM
ns
**
LDH release %
80
60
40
20
0
Untreated
LPS/Nig/scRNA
LPS/Nig/ZDHHC5 siRNA
LPS/Nig/ZDHHC9 siRNA
d
mBMDM
- + + LPS/Nig
- + - scRNA
(kDa) - - + ZDHHC5 SiRNA
52 GSDMD
30 GSDMD-NT
71 ZDHHC5
45 Actin
mBMDM
- + + LPS/Nig
- + - scRNA
(kDa) - - + ZDHHC9 SiRNA
52 GSDMD
30 GSDMD-NT
45 ZDHHC9
45 Actin

**Fig. S10.**

**(a) Expression of palmitoylation-related genes in THP-1 cells.** Real-time PCR analysis of THP-1 cells primed with LPS for 3 h. mRNA expression of ZDHHC was quantified and normalized to actin. Data are mean ± SEM. *P<0.05, **P<0.01, ***P<0.001, unpaired two-tailed Student's *t*-test.

**(b-d) ZDHHC5 palmitoyl enzyme mediates pyroptotic cell death in primary mouse macrophages.**

(c) Fluorescence images of SYTOX Green uptake in primary mouse bone marrow-derived macrophages transfected with control or ZDHHC siRNA. Right, image represents the quantification of SYTOX uptake in mBMDMs. Experiments were performed three times. Data are mean ± SEM. *P<0.05, unpaired two-tailed Student's *t*-test.

(c) LDH release in cell culture supernatants from mBMDM cultures was quantified. Data are mean ± SEM. **P<0.01, unpaired two-tailed Student's *t*-test.

(d) mBMDMs transfected with control or *Zdhhc* siRNAs were treated with or without LPS/Nig. Cells were lysed and subjected to western blot analysis. Western blot images are representative of three independent repeats.

Fig.S11

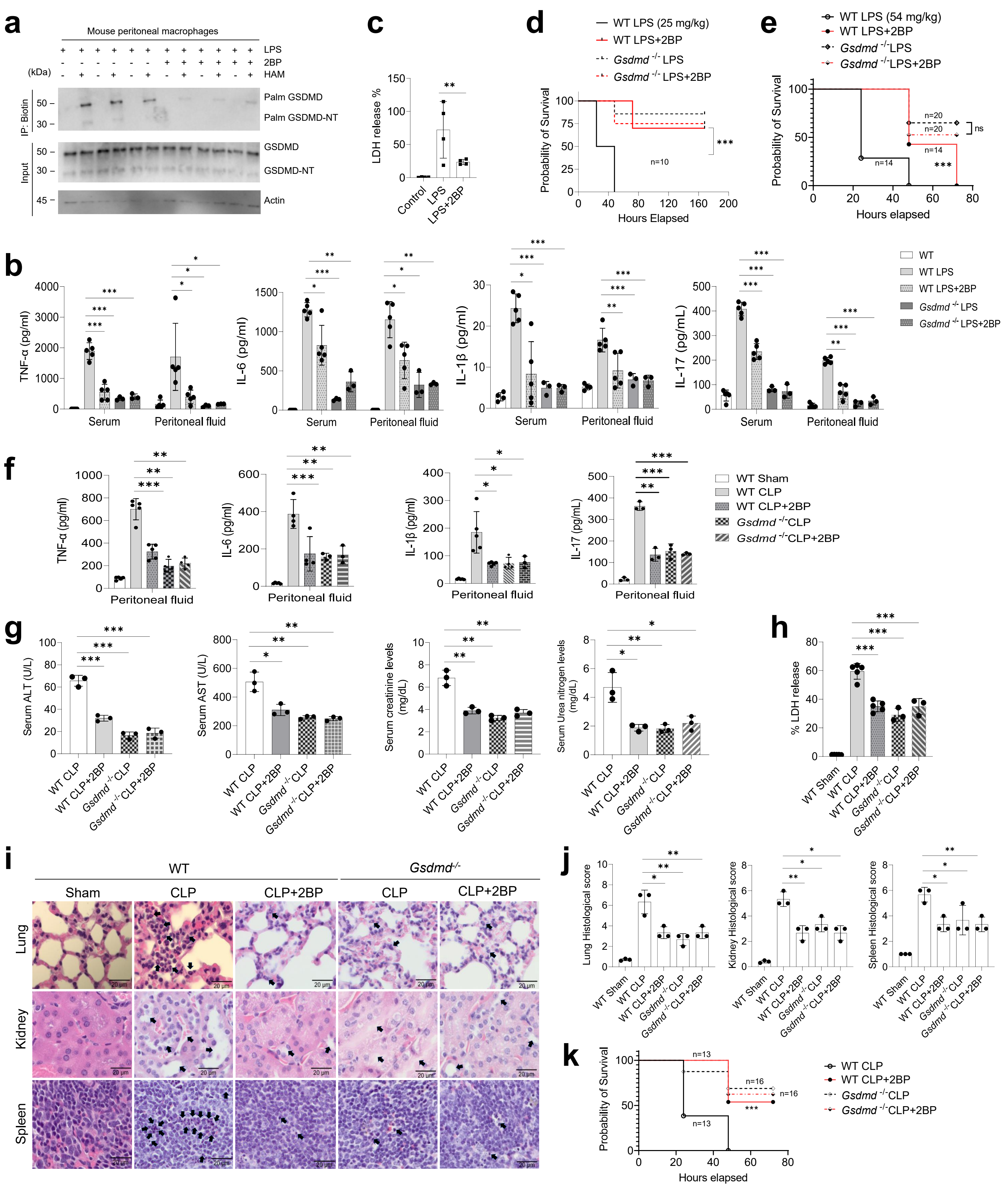
a
Mouse peritoneal macrophages
LPS
2BP
HAM
(kDa)
IP: Biotin
Palm GSDMD
Palm GSDMD-NT
Input
GSDMD
GSDMD-NT
Actin
c
LDH release %
Control
LPS
LPS+2BP
d
WT LPS (25 mg/kg)
WT LPS+2BP
Gsdmd -/- LPS
Gsdmd -/- LPS+2BP
Probability of Survival
Hours Elapsed
n=10
e
WT LPS (54 mg/kg)
WT LPS+2BP
Gsdmd -/-LPS
Gsdmd -/-LPS+2BP
Probability of Survival
Hours elapsed
ns
b
TNF-α (pg/ml)
IL-6 (pg/ml)
IL-1β (pg/ml)
IL-17 (pg/mL)
Serum
Peritoneal fluid
WT
WT LPS
WT LPS+2BP
Gsdmd -/- LPS
Gsdmd -/- LPS+2BP
f
WT Sham
WT CLP
WT CLP+2BP
Gsdmd -/-CLP
Gsdmd -/-CLP+2BP
g
Serum ALT (U/L)
Serum AST (U/L)
Serum creatinine levels (mg/dL)
Serum Urea nitrogen levels (mg/dL)
h
% LDH release
i
WT
Gsdmd-/-
Sham
CLP
CLP+2BP
Lung
Kidney
Spleen
20 μm
j
Lung Histological score
Kidney Histological score
Spleen Histological score
k
Probability of Survival
Hours elapsed
WT CLP
WT CLP+2BP
Gsdmd -/-CLP
Gsdmd -/-CLP+2BP

**Fig. S11. 2-bromopalmitate (2BP), a palmitoylation inhibitor, alleviates inflammation and prolongs mouse survival in sepsis.**

(a) Peritoneal mouse macrophages were harvested from the indicated groups of mice either pretreated or not with 2BP (30 mg/kg) for 6 h followed by intraperitoneal challenge or not with 20 mg/kg LPS for 12 h. Peritoneal cells were lysed and blocked with NEM and subjected to the ABE palmitoylation assay in the presence or absence of HAM. Immunoblot analyses were performed with anti-GSDMD antibodies. Data are representative of three independent experiments.

(b) Shown are quantities of IL1-β, TNF-α, IL-17, and IL-6 present in the serum and peritoneal fluid of LPS-induced septic wild type and *Gsdmd*$^{-/-}$ mice. n=5 mice per group. *P<0.05, **P<0.01, ***P<0.001, unpaired Student's *t*-test.

(c) The peritoneal fluid was collected and centrifuged at 16,000 x g for 10 min before quantification of LDH release. n=4 mice per group. *P<0.05, **P<0.01, ***P<0.001, unpaired Student's *t*-test.

(d) WT or *Gsdmd* KO mice were pretreated with 2BP (30 mg/kg) or vehicle (Ctrl) by IP injection 4 h before intraperitoneal challenge with 25 mg/kg LPS and monitoring for survival. Survival rates were analyzed using Kaplan-Meier survival curves and log-rank testing. n=10 mice per group. *P<0.05, **P<0.01, ***P<0.001.

(e) *Gsdmd*$^{-/-}$ mice were pre-treated with vehicle or 2BP (30 mg/kg) for 4 h followed by LPS treatment (54 mg/kg) and monitoring for survival. Survival rates were calculated using Kaplan-Meier survival curves. *P<0.05, **P<0.01, ***P<0.001.

(f) Peritoneal fluid IL1-β, TNF-α, IL-17, and IL-6 levels were measured from CLP-induced sepsis mice followed by treatment with or without 2BP by ELISA. n≥3 mice per group. *P<0.05, **P<0.01, ***P<0.001, unpaired Student's *t*-test.

(g) Shown are circulating AST, ALT, BUN, and creatine levels in the serum of CLP-induced wild type and *Gsdmd*$^{-/-}$ mice. n=3 mice per group. *P<0.05, **P<0.01, ***P<0.001, unpaired Student's *t*-test. ALT: alanine transaminase, AST: aspartate aminotransferase, BUN: blood urea nitrogen.

(h) LDH release was quantified from the peritoneal fluid of CLP-induced sepsis mice. n=5 mice per group. *P<0.05, **P<0.01, ***P<0.001, unpaired Student's *t*-test.

(i) Histopathologic assessment of WT and *Gsdmd*$^{-/-}$ mice one day after CLP-induced sepsis followed by treatment with or without 2BP (30 mg/kg). Shown are representative H&E-stained sections of kidney, liver, and lung tissues from three independent experiments.

(j) Histopathological scores. Student's *t*-test was used to compare the means between two groups. * indicates p<0.05, ** indicates p<0.01. Data presented are means ± SD (n=3).

(k) Survival analyses of CLP-induced sepsis mice treated with or without 2BP (30 mg/kg). n=5 mice per group. Survival rates were calculated using Kaplan-Meier survival curves. *P<0.05, **P<0.01, ***P<0.001, log-rank test.

Fig.S12

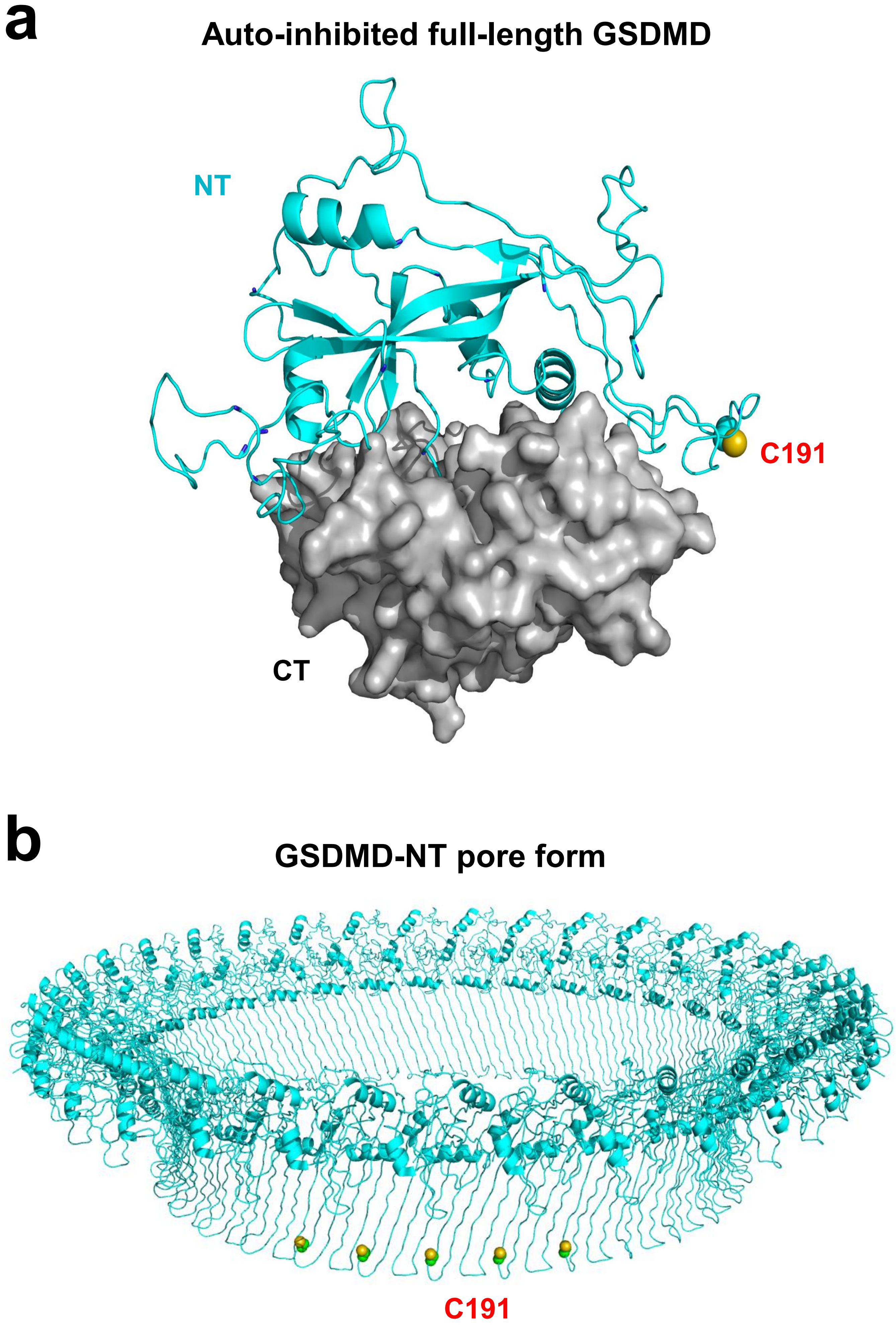
a
Auto-inhibited full-length GSDMD
NT
C191
CT
b
GSDMD-NT pore form
C191

**Fig. S12. Cys191 in auto-inhibited full-length GSDMD and GSDMD-NT pore structures.**

(a) The available crystal structure of GSDMD (PDB: 6n9o) does not contain the C191 region (*34*). Homology model of full-length human GSDMD generated by SWISS-Model (*35*) based on 6n9o revealed the potentially exposed C191 residue. The side chain of C191 is shown in atomic spheres with the sulfur atom in gold and carbon atoms in cyan.

(b) The cryo-EM structure of human GSDMD-NT pore with 33-fold symmetry (PDB: 6vfe) is displayed here with the C191 residue side chain for five consecutive subunits highlighted as atomic spheres. The sulfur atom is colored gold and carbon atoms in cyan. Electron microscopy imaging, data processing, model building, and structure analysis were conducted as previously described (*36*).

**Supplementary Table S1.**

Excel file depicts the proteins identified in the GSDMD-Flag, GFP-FLAG, actin, and IgG alone samples (**Fig. 1a)** using mass spectrometry. Data analysis was performed using Proteome Discoverer (Thermo Fisher Scientific). Raw mass spectrometric data were deposited in ProteomeXchange Consortium with the dataset identifier PXD039133.

**Supplementary Table S2.**

Protein hits identified in the GSDMD-FL sample by mass spectrometry classified according to GO analyses (biological process, cellular component, and molecular function) using the Metascape tool (metascape-org).
