## Supplementary Table 2 for "Palmitoylation of gasdermin D directs its membrane translocation and pore formation in pyroptosis"

| **Gene** | **Protein name** | **Biological process** | **Cellular component** | **Molecular function** |
| --- | --- | --- | --- | --- |
| **GSDMD** | **gasdermin D** | **pore formation in membrane** | **NLRP3 inflammasome complex** | **phosphatidic acid binding** |
| HSP90AB1 | heat shock protein 90 alpha family class B member 1 | negative regulation of transforming growth factor beta activation | HSP90-CDC37 chaperone complex | UTP binding |
| EEF2 | eukaryotic translation elongation factor 2 | translational elongation | aggresome | translation elongation factor activity |
| PKM | pyruvate kinase M1/2 | positive regulation of cytoplasmic translation | rough endoplasmic reticulum | pyruvate kinase activity |
| RIOK1 | RIO kinase 1 | positive regulation of rRNA processing | preribosome | protein serine kinase activity |
| PPM1B | protein phosphatase, Mg2+/Mn2+ dependent 1B | N-terminal protein myristoylation | nucleolus | manganese ion binding |
| BCAP31 | B cell receptor associated protein 31 | positive regulation of retrograde protein transport | mitochondria-associated endoplasmic reticulum membrane | MHC class I protein binding |
| HSP90AA1 | heat shock protein 90 alpha family class A member 1 | protein unfolding | dendritic growth cone | TPR domain binding |
| WDR77 | WD repeat domain 77 | histone H4-R3 methylation | methylosome | methyl-CpG binding |
| SLC16A1 | solute carrier family 16 member 1 | plasma membrane lactate transport | lateral plasma membrane | mevalonate transmembrane transporter activity |
| ATP2A2 | ATPase sarcoplasmic/endoplasmic reticulum Ca2+ transporting 2 | calcium ion transport from cytosol to endoplasmic reticulum | platelet dense tubular network membrane | S100 protein binding |
| EEF1A1 | eukaryotic translation elongation factor 1 alpha 1 | translational elongation | eukaryotic translation elongation factor 1 complex | translation elongation factor activity |
| MAP1B | microtubule associated protein 1B | regulation of synaptic plasticity by chemical substance | hippocampal mossy fiberr | microtubule binding |
| PHGDH | phosphoglycerate dehydrogenase | gamma-aminobutyric acid metabolic process | extracellular exosome | phosphoglycerate dehydrogenase activity |
| LBR | lamin B receptor | neutrophil differentiation | integral component of nuclear inner membrane | delta14-sterol reductase activity |
| PRMT5 | protein arginine methyltransferase 5 | regulation of adenylate cyclase-inhibiting dopamine receptor signaling pathway methylation | methylosome | histone methyltransferase activity (H4-R3 specific) |
| UQCRC2 | ubiquinol-cytochrome c reductase core protein 2 | mitochondrial electron transport | mitochondrial respiratory chain complex III | metalloendopeptidase activity |
| CHCHD3 | coiled-coil-helix-coiled-coil-helix domain containing 3 | cristae formation | mitochondrial crista junction | phosphatase binding |
| EIF4B | eukaryotic translation initiation factor 4B | eukaryotic translation initiation factor 4F complex assembly | eukaryotic translation initiation factor 4F complex | RNA strand-exchange activity |
| RPN1 | ribophorin I | protein N-linked glycosylation via asparagine | oligosaccharyltransferase complex | dolichyl-diphosphooligosaccharide-protein glycotransferase activity |
| **FASN** | **fatty acid synthase** | **Lipid biosynthetic process** | **glycogen granule** | **[acyl-carrier-protein] S-acetyltransferase activity** |
| CCT3 | chaperonin containing TCP1 subunit 3 | positive regulation of establishment of protein localization to telomere | zona pellucida receptor complex | protein folding chaperone |
| STK38 | serine/threonine kinase 38 | negative regulation of MAP kinase activity | cytosol | mitogen-activated protein kinase kinase kinase binding |
| NDUFS1 | NADH:ubiquinone oxidoreductase core subunit S1 | mitochondrial electron transport | mitochondrial respiratory chain complex I | 2 iron, 2 sulfur cluster binding |
| IRS4 | insulin receptor substrate 4 | insulin receptor signaling pathway | cytosol | insulin receptor binding |
| EMD | emerin | positive regulation of protein export from nucleus | cortical endoplasmic reticulum | beta-tubulin binding |
| ACSL3 | acyl-CoA synthetase long chain family member 3 | positive regulation of phosphatidylcholine metabolic process | peroxisomal membrane | palmitoyl-CoA ligase activity |
| RPS2 | ribosomal protein S2 | positive regulation of ubiquitin-protein transferase activity | cytosolic small ribosomal subunit | fibroblast growth factor binding |
| SLC25A3 | solute carrier family 25 member 3 | mitochondrial phosphate ion transmembrane transport | integral component of mitochondrial inner membrane | phosphate:proton symporter activity |
| GCN1 | GCN1 activator of EIF2AK4 | cellular response to leucine starvation | polysome | ribosome binding |
| RPS11 | ribosomal protein S11 | cytoplasmic translation | cytosolic small ribosomal subunit | rRNA binding |
| SPIN1 | spindlin 1 | rRNA transcription | nuclear membrane | methylated histone binding |
| CCT8 | chaperonin containing TCP1 subunit 8 | positive regulation of establishment of protein localization to telomere | zona pellucida receptor complex | protein folding chaperone |
| SMC2 | structural maintenance of chromosomes 2 | positive regulation of chromosome condensation | condensin complex | single-stranded DNA binding |
| CALU | calumenin | GO:0008150 biological_process | sarcoplasmic reticulum lumen | calcium ion binding |
| MYH10 | myosin heavy chain 10 | mitotic cytokinesis | myosin II filament | RNA stem-loop binding |
| SMC4 | structural maintenance of chromosomes 4 | positive regulation of chromosome condensation | condensin complex | single-stranded DNA binding |
| AIFM1 | apoptosis inducing factor mitochondria associated 1 | protein import into mitochondrial intermembrane space | mitochondrial intermembrane space | NAD(P)H oxidase H2O2-forming activity |
| NNT | nicotinamide nucleotide transhydrogenase | NADPH regeneration | mitochondrial respirasome | NAD(P)+ transhydrogenase (B-specific) activity |
| CCT2 | chaperonin containing TCP1 subunit 2 | chaperone mediated protein folding independent of cofactor | chaperone complex | protein folding chaperone |
| RPS8 | ribosomal protein S8 | ribosomal small subunit biogenesis | cytosolic small ribosomal subunit | structural constituent of ribosome |
| CTTN | cortactin | substrate-dependent cell migration, cell extension | mitotic spindle midzone | profilin binding |
| RCN2 | reticulocalbin 2 | ER component | endoplasmic reticulum lumen | calcium ion binding |
| ATP5F1C | ATP synthase F1 subunit gamma | proton motive force-driven mitochondrial ATP synthesis | mitochondrial proton-transporting ATP synthase complex, catalytic sector F(1) complex, catalytic domain | proton-transporting ATP synthase activity |
| RPL28 | ribosomal protein L28 | cytoplasmic translation | cytosolic large ribosomal subunit | structural constituent of ribosome |
| PRKDC | protein kinase, DNA-activated, catalytic subunit | regulation of platelet formation | DNA-dependent protein kinase complex | DNA-dependent protein kinase activity |
| RIF1 | replication timing regulatory factor 1 | subtelomeric heterochromatin assembly | chromosome, telomeric repeat region | protein binding |
